# CANDOCK: Chemical atomic network based hierarchical flexible docking algorithm using generalized statistical potentials

**DOI:** 10.1101/442897

**Authors:** Jonathan Fine, Janez Konc, Ram Samudrala, Gaurav Chopra

## Abstract

Small molecule docking has proven to be invaluable for drug design and discovery. However, existing docking methods have several limitations, such as, improper treatment of the interactions of essential components in the chemical environment of the binding pocket (e.g. cofactors, metal-ions, *etc.*), incomplete sampling of chemically relevant ligand conformational space, and the inability to consistently correlate docking scores of the best binding pose with experimental binding affinities. We present CANDOCK, a novel docking algorithm that utilizes a hierarchical approach to reconstruct ligands from an atomic grid using graph theory and generalized statistical potential functions to sample biologically relevant ligand conformations. Our algorithm accounts for protein flexibility, solvent, metal ions and cofactors interactions in the binding pocket that are traditionally ignored by current methods. We evaluate the algorithm on the PDBbind and Astex proteins to show its ability to reproduce the binding mode of the ligands that is independent of the initial ligand conformation in these benchmarks. Finally, we identify the best selector and ranker potential functions, such that, the statistical score of best selected docked pose correlates with the experimental binding affinities of the ligands for any given protein target. Our results indicate that CANDOCK is a generalized flexible docking method that addresses several limitations of current docking methods by considering all interactions in the chemical environment of a binding pocket for correlating the best docked pose with biological activity.

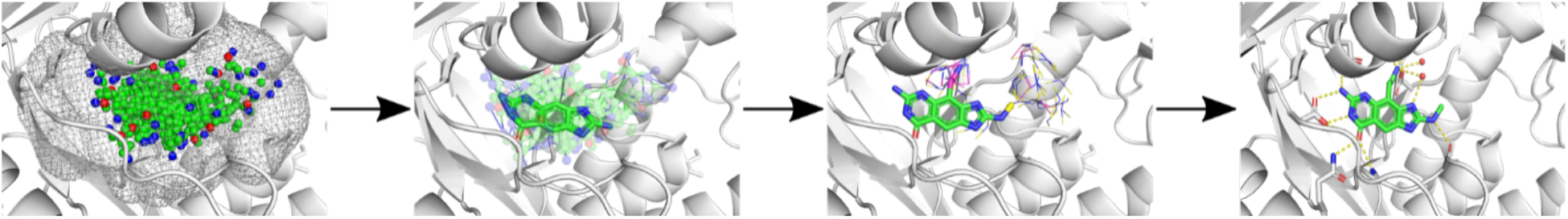

## 1. Introduction

Computational docking provides a means to predict and assess interactions between ligands and proteins with relatively little investment. Docking refers to physical three-dimensional structural interactions between a receptor (typically, proteins, DNA, RNA, *etc*.) and a ligand (small molecules, proteins, peptides, *etc*.)^1–15^. Docking methods are evaluated by predicting the correct pose/binding mode (evaluated using RMSD or TMScore of the coordinates of the atoms) or by measuring predicted binding affinities^4,8,11,12,16^. Application to protein targets involved in disease holds the promise of discovering new therapeutics using traditional single target approaches or by virtually measuring the interactions of a compound with the proteins from multi-organism proteome^17–22^. The resulting chemo-proteome interactions can be interrogated to study polypharmacology^19^ and investigate the effect drugs and agents have on protein classes in a disease-specific context^19,22^. In previous works, we have used the algorithm presented herein to combat Ebola^20^, determine the toxicity of potential diabetes therapeutics^21^, and rank the affinity of kinase inhibitors for the treatment of Acute Myeloid Leukemia^23^.

More than 20 molecular docking software tools, such as, Autodock Vina^24^, Gold^25^, and Glide^3^, are currently in use for pharmaceutical research. However, after decades of method development and application, the promise to computationally determine new therapeutics has not been fully realized and computational methods for drug discovery are still in its infancy^26,27^. The CANDOCK algorithm confronts several outstanding technical and practical problems in computational docking. For example, one significant problem is assessing goodness-of-fit, or the likelihood that the given pose is the most physically realistic (native-like) pose among many unrealistic binding poses. Another significant limitation is the lack of full protein flexibility in the docking methods used today. The induced fit is a widely recognized challenge in computational drug screening^28^, where the protein and the ligand undergo conformational changes upon ligand binding. Therefore, the traditional treatment of proteins as rigid structures may be insufficient and often misleading for structure-guided drug screening and design as shown by us and others previously^29^. Docking ligands to their protein targets is particularly challenging when attempting to reproduce the binding mode of small molecules to ligand-free or alternative ligand-bound protein structures, which invariably occurs for practical application of any docking method. Specifically, docking with ligand-bound (holo) protein structures typically leads to an accuracy of 60-80%, whereas ligand-free (apo) structures yields a docking accuracy of merely 20-40%^30–34^.

Several methods have been implemented to account for protein and ligand flexibility, including multiple experimentally derived structures from X-ray crystallography^35^, nuclear magnetic resonance^35^ rotamer libraries^36,37^, Monte Carlo^24,38^, and molecular mechanics^39–44^. The same principle limits use of multiple experimentally derived protein structures or side-chain rotamer libraries: binding a ligand to a protein can cause conformational changes in either molecule that are not captured by these methods^45^. The sampling problem is compounded by the fact that the protein main chain torsion angles are also frequently altered from their ligand-free conformations, which these methods fail to capture. Molecular mechanics is well suited for capturing fine detail side-chain and main chain motions and rearrangements through energy minimization. However, molecular mechanics is limited in that adequate sampling of all degrees of freedom between protein and ligand: rotation, translation, and torsion angle are frequently computationally intractable. Further, the use of unrestrained molecular dynamics has been shown to disrupt the ligand from its native pose^46^.

Modern docking methods address these issues by employing algorithms such as the Genetic Algorithm^25,28,47,48^ to flexibly sample the conformational space. However, it has been shown that these methods do not consistently produce poses that rank the biological activity of the ligand well^48,49^ and that the ability of these methods to produce a correct pose is dependent on the starting conformation of the ligand^50,51^. Some methodologies use a fragment-based approach to docking^52^ to sample the conformational space for a given ligand efficiently. These fragment-based methods have reported a greater ability to rank activity between given ligands^53,54^. Therefore, we believe that further innovation in fragment-based methods is an appropriate way to improve docking methods.

We have developed the CANDOCK algorithm around a new protocol for hierarchical (atoms to fragments to molecules) docking with iterative dynamics during molecule reconstruction to “grow” the ligand in the binding pocket. The docking protocol is based on two guiding principles: (i) binding sites possess regions of both very high and very low structural stability^55^ and (ii) a tandem sequence of small protein motions are generally sufficient to predict the correct binding mode of protein-ligand interactions^45^. The hierarchical nature of this method is derived from an ‘atoms to fragments,’ ‘fragments to ligands’ approach that generates chemically relevant poses given the ligand and surrounding any chemical environment (e.g. protein, RNA, DNA binding sites or interfaces). For any flexible ligand, the expectation is that at least one or a few fragments conformations assembled using ligand-receptor atomic interactions in the binding pocket will bind to a structurally stable region of the receptor. Following identification of such a binding mode, subtle conformational changes of the receptor is necessary for reconstructing the ligand using these fragments as “seeds” to generate accurate receptor-ligand binding modes (poses). We show that CANDOCK can accurately reproduce the binding mode of ligands and rank the activity of these ligands in such poses using a generalized statistically derived forcefield, demonstrating the potential to overcome traditional challenges with induced-fit docking methods.

## 2. Materials and methods

We first introduce our generalized statistical scoring function, then provide details of the CANDOCK algorithm, and selection of benchmarking datasets for evaluating pose selection and receptor-ligand affinity ranking.

### 2.1. Generalized statistical scoring function

A generalized statistical scoring potential is used to account for varying chemical environments, such as metal ions, cofactors, water molecules, *etc*. The scoring function employed by the CANDOCK algorithm is a pairwise atomic scoring function that is based on our previous work^56^. Here, we reproduce the fundamental equations^56^ to clarify the terminology used in our manuscript. The scoring function calculates the potential between two atoms based on the distance between atoms *i* and *j* with atom types *a* and *b and* takes four input terms that determine the method by which score is calculated. The possible terms are ‘functional’, ‘reference’, ‘composition’, and ‘cutoff’ which define the probability function *P* given in Eq. (1):

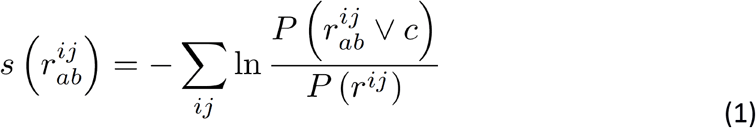

The ‘functional’ term determines the numerator of Eq. (1) and can be defined either as a ‘normalized frequency’ function *f(r)* in Eq. (2) or a ‘radial’ distribution function *g(r)* given in Eq. (3):

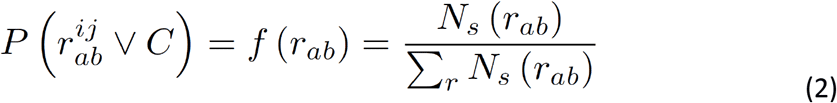

where *Ns* is the number of observed atoms found at a given distance.

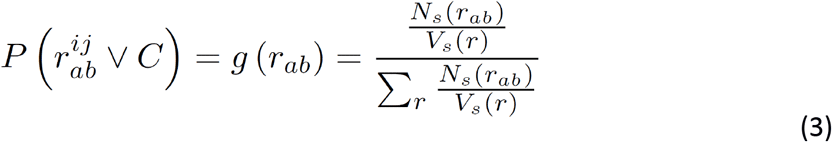

where *N*_*s*_ is divided by the volume of the sphere *V*_*s*_(*r*). To distinguish between these two functions, ‘radial’ scoring functions start with ‘R’ while ‘normalized frequency’ functions start with ‘F’. The ‘reference’ term determines the denominator of the scoring function. It can be defined either as ‘mean’, in which case it is calculated as a sum of all atom type pairs divided by the number of atom types. This term can be used with either ‘normalized frequency’ (Eq. (4)) or ‘radial’ (Eq. (5)):

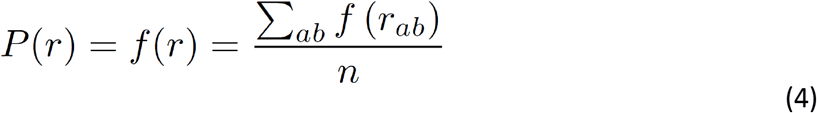

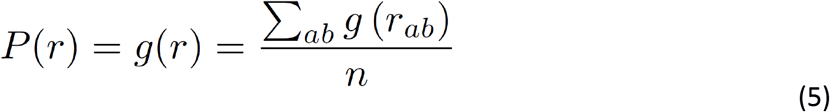

The second option is the ‘cumulative’ which denotes cumulative distribution. Used together with ‘normalized frequency’ this yields Eq. (6) and ‘radial’ yields Eq. (7):

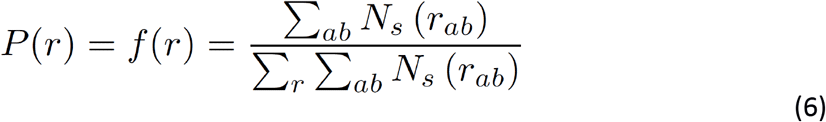

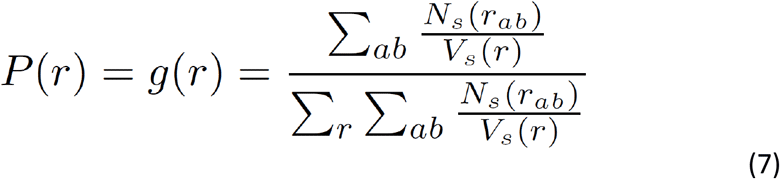

Scoring functions compiled with the ‘mean’ option are denoted as ‘M’ while those compiled with the ‘cumulative’ are denoted as ‘C’. The third term defines the composition of the scoring function. This term controls the number of unique atom pairs used for compiling the scoring function. The ‘complete’ option will result in the scoring function compiled from all possible atom type pairs while the ‘reduced’ option will cause only atom pairs present in the given complex to be used. The letter ‘C’ is used to denote complete scoring function while ‘R’ is used to denoted scoring function that is compiled with the ‘reduced’ option. A total of 8 scoring function families can be created with these three options (RMR, RMC, RCR, RCC, FMR, FMC, FCR, FCC). The fourth and final term used to compile the scoring function is the ‘cutoff’ which controls the maximum distance at which the interactions will be calculated, with possible values ranging from 4 Å to 15 Å. With all four options there are a total of 96 possible scoring functions (8×12) to account for generalized parameters for identifying native poses and activity across a diverse set of biomolecular interactions in varying chemical environments (proteins, nucleic acids, interfaces, cofactors, etc.). Example scoring functions are, ‘radial-mean-reduced-6’ (RMR6), ‘normalized frequency-cumulative-complete-8’ (FCC8), etc. as denoted in the manuscript.

### 2.2. The CANDOCK algorithm

#### 2.2.1. Phase I: Structure Preparation

The CANDOCK algorithm’s input is a set of compounds to be docked, a query protein structure, and a set of binding sites on the query protein structure. In a three-phase protocol (Figure 1), it performs semi or fully flexible docking of compounds to the protein and outputs docked and minimized protein-compound complex structures together with their predicted scores.

**Figure 1:**
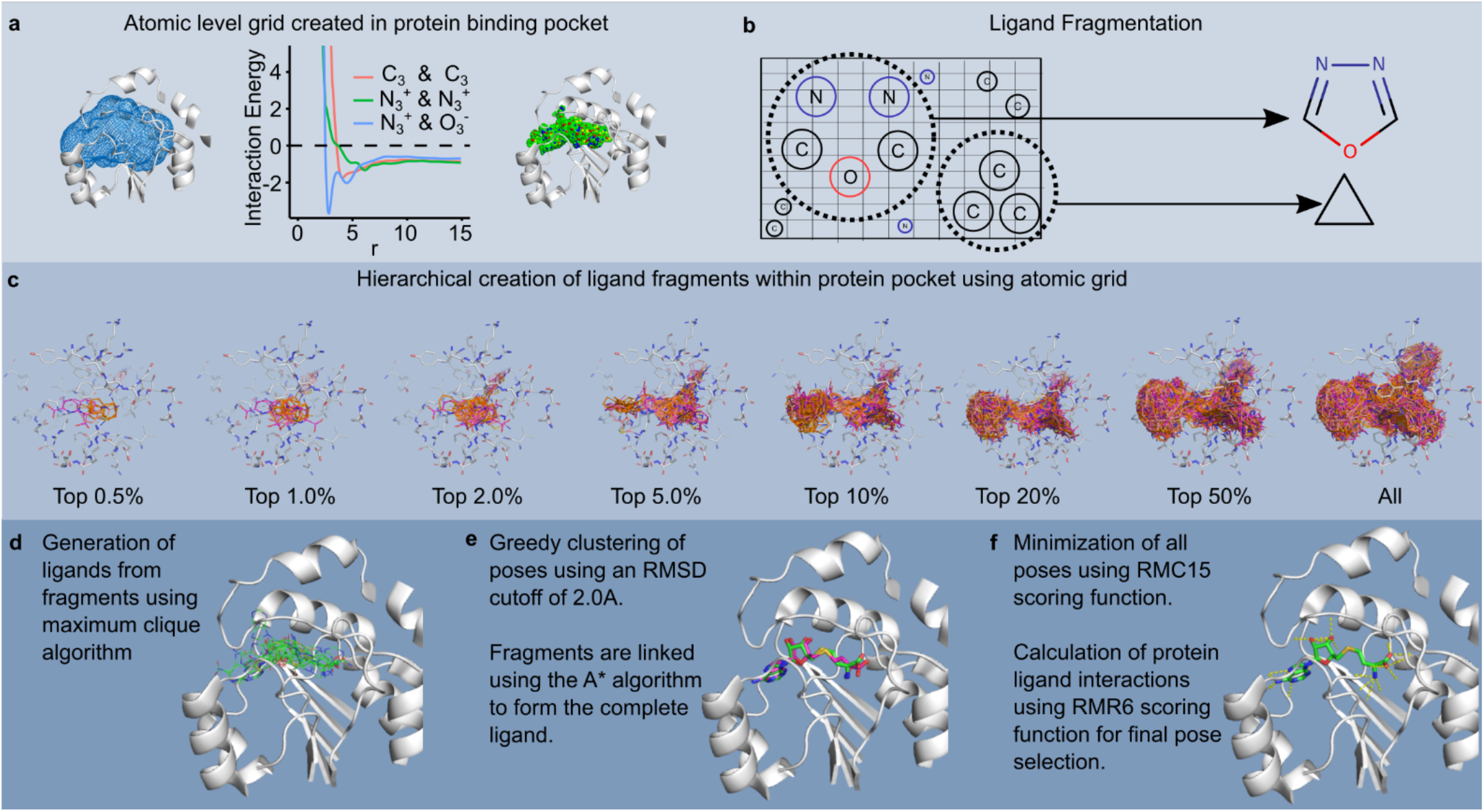
Overview of the CANDOCK docking algorithm. Phase I consists of processing the input protein (**a**) and the ligand (**b**). During Phase I, an atomic grid is created in the protein binding site where the scores of all possible atom types at each point in the binding site grid. Simultaneously, the input ligand(s) are fragmented along the rotatable bonds present in the ligand. The grid is used to recreate the rigid fragments in the binding pocket. Phase II constructs the rigid ligand fragments in the binding site grid producing ‘seeds’ that can be grown into the full ligand (**c**). Phase III identifies potential ligand poses using maximum clique algorithm (**d**), clusters and links these poses using A*(**e**) and minimizes the poses into the binding site (**f**).

##### Parse receptor and compounds

The inputs to the algorithm are the 3D coordinates and topology of a query receptor (e.g. protein structure) consisting of single or multiple chains which may also contain cofactors and post-translation modifications in the PDB format, and compounds in the MOL2 format. Compounds are processed in batches of size 10 to enable reading of large molecular files that do not fit in computer memory. An example of a ligand is given in Figure 2a.

**Figure 2:**
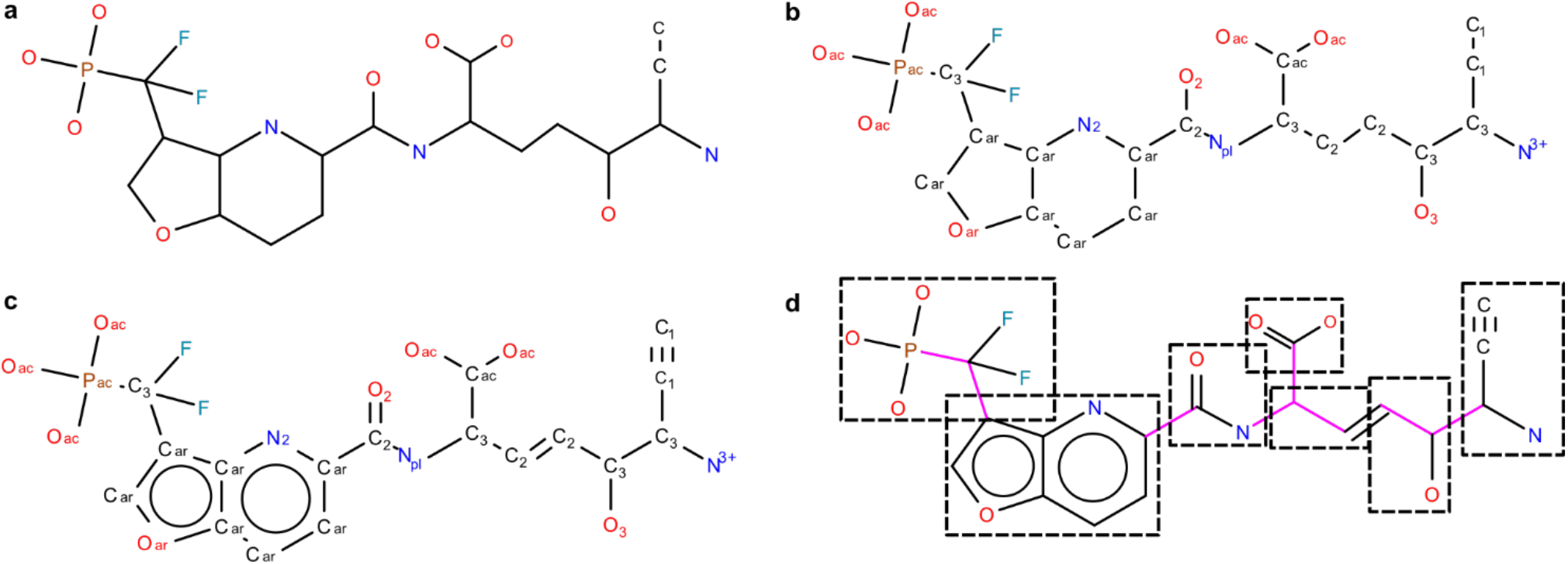
Atom type assignment and fragmentation procedure present in CANDOCK. The procedure begins with the topology and 3D coordinates of the ligand **(a)**. Using these data, the IDATM type is assigned to each atom in the ligand using a previously described algorithm^27^). This yields the hybridization state of all atoms, allowing for the assignment of bond orders for all atoms **(c)**. The bond orders and topologies are used to assign a rotatable flag for each bond in the ligand using rules derived from the DOCK 6 program^31^. The rigid fragments identified using this method are boxed **(d)**.

##### Compute atom types

To compute atom types for protein, cofactors, and compounds, we implemented the IDATM algorithm^57^ (results given in Figure 2b). We also implemented an algorithm^58,59^ to assign AMBER General Force Field (GAFF) atom types to cofactors, ligands, and post-translational modifications, while GAFF types for proteins are obtained from the AMBER10 topology file available as part of the OpenMM package^60^.

##### Assignment of bond orders

Using the hybridization information provided by the newly assigned IDATM atom types, several potential bond order states can be generated as to fit with the expected number of bonds (valence) for each ligand atom. These potential bond order assignments are evaluated in a trial and error fashion to determine whether they form a valid molecule using valence state rules derived for all atom types. The bond order set that satisfies the set of valence states with the lowest sum of atomic penalty scores over all atoms (see Figure 2c) is used to assign GAFF bond orders of the ligand.

##### Fragment compounds

Rotatable bonds are first identified in each compound using the extended list of rotatable bonds adapted from the UCSF DOCK 6 software^61^. Next, structurally rigid fragments consisting of atoms between the rotatable bonds are identified. Bond vectors for rotatable bonds are retained for each rigid fragment to be used during reconstruction of docked fragments. Fragments consisting of more than 4 atoms, in which at least two atoms are rigid (connected by a non-rotatable bond) are considered as seed fragments. These are subsequently rigidly docked into the protein binding site. All other non-seed fragments are considered as linking fragments during the compound reconstruction process. This result is shown in Figure 2d.

##### Assignment of force field atom types

Using the computed GAFF atom types, the bonded forces of the AMBER force field are generated for the protein and the docked compounds. Protein-compound interactions are scored using the knowledge-based Radial Mean Reduced (RMR) discriminatory function defined previously^56^ with a 6 Å cutoff (see section on Generalized statistical scoring function). This function calculates a fitness score for each compound’s or fragment’s atom in a protein by considering all protein atoms within 6 Å radius of that atom. It is an atomic level radial distribution function with mean reference state that averages over all pairwise atom types from a reduced atom type composition (protein’s and compound’s atom types), using experimentally determined intermolecular complexes in the Cambridge Structural Database (CSD)^62^ and in the Protein Data Bank (PDB)^63^ as the information sources. The objective function that is used for the minimization of the protein-compound interactions is computed using the RMC scoring function with a 15 Å cutoff as follows: for each possible pair of atom types present in the protein-ligand complex, the RMC function is sampled at discrete 0.1 Å intervals and is smoothed using B-spline interpolation. Potential energy values and their first derivatives are calculated at 0.01 Å intervals over the [0, 15] Å interval for the smoothed function. The objective function is implemented as a custom knowledge-based force object in OpenMM^60^ which is used as a library from the CANDOCK source code.

##### Prepare protein for molecular mechanics

The N- and C-terminal residues are renamed according to the AMBER topology specification, e.g., ALA to NALA or CALA, disulfide bonds are added to the protein by connection of SG atoms that are closer than 2.5 Å, inter-residue bonds are also added by connection of main chain C and N atoms that are closer than 1.4 Å.

#### 2.2.2. Phase II: Rigid Fragment Docking

##### Compute rotations of seeds

For each seed fragment, we compute its rotational transformations about the geometric center which is fixed at the coordinate origin. Accordingly, we first compute 256 uniformly distributed unit vectors around the coordinate origin. Then, the seed fragment is rotated by 10° increments around the axis formed by each unit vector. To speed up the subsequent step of rigid fragment docking, the rotated fragment atoms’ coordinates are mapped on a hexagonal close-packed (HCP) grid of 0.375 Å resolution. This mapping enables efficient docking of fragments to a protein binding site since their rotational transformations need to be computed only once. The fragment’s clashes with the protein and the fragment’s RMR6 scores are determined by translations of the rotational fragment grid over the compatible HCP binding site grid using fast integer arithmetic.

##### Generate binding site grid

A binding site location for docking is specified using one or more centroids, each consisting of the Cartesian coordinate of its center and its radius. We generate a grid that covers the space of all centroids that represent the binding site (Figure 3a). We use an HCP grid that provides maximal packing efficiency, covering the same volumetric space of a simple cubic grid with approximately 40% fewer grid points to achieve the same maximal interstitial spacing. The grid points are in a distance range of 0.8 Å < d < 8 Å from any protein atom. We use a grid spacing of 0.375 Å with a maximal interstitial spacing of 0.22 Å to densely represent the protein binding sites (Figure 3b).

**Figure 3:**
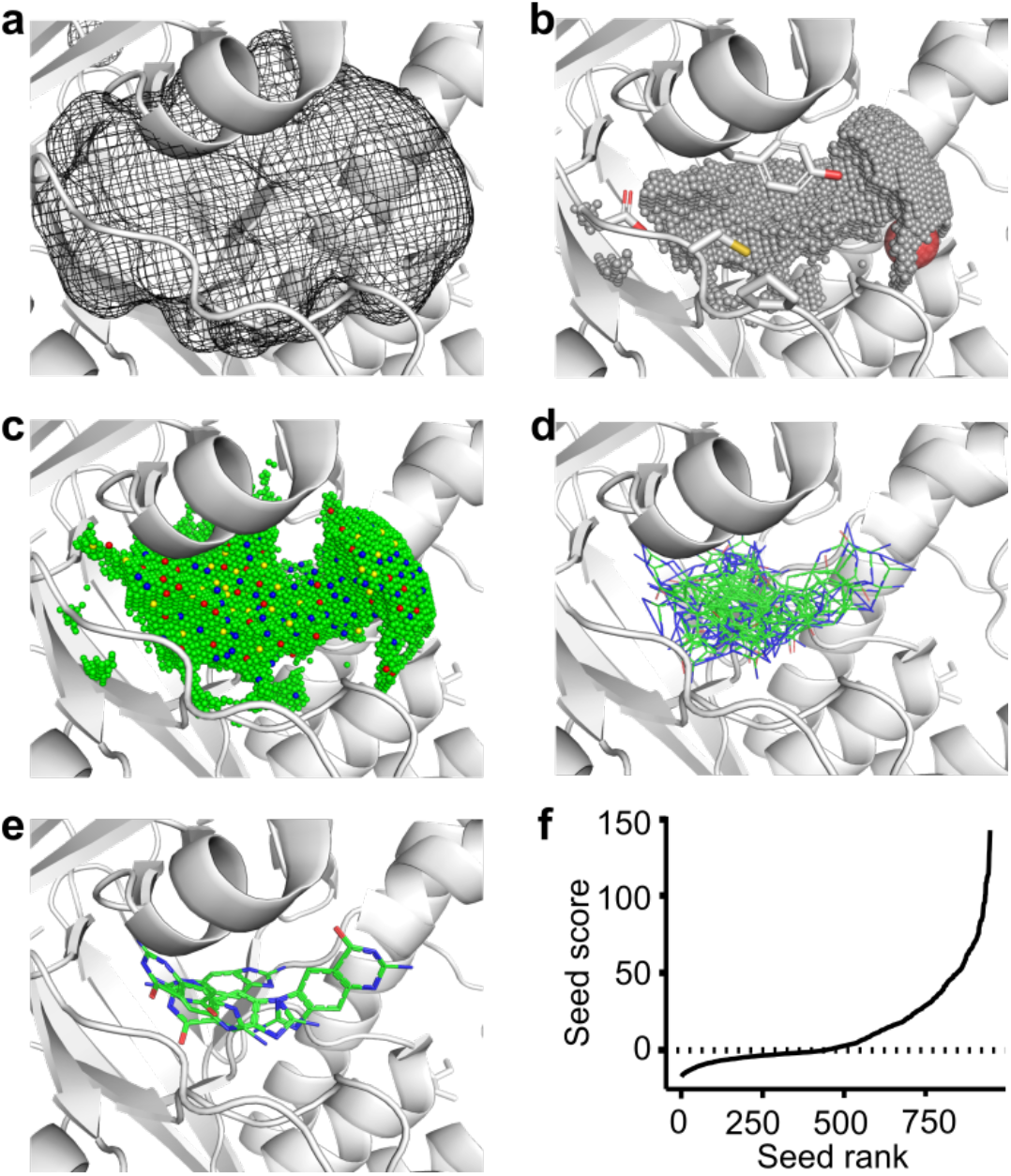
Detailed overview of the hierarchal relationship between the atomic grid and ligand fragments. The protein binding site is supplied as a series of centroids that are combined to form a volume of space that defines the binding pocket **(a)**. Regions of this volume that do not clash with the protein, waters, or cofactors are filled with a hexagonal close-packed grid **(b)**. The score of all atom types present in the ligand are calculated at each grid point using the RMR6 scoring function **(c)**. Ligand fragments from the previous step are translated and rotated within this grid to produce a collection of the same ligand fragment throughout the binding site **(d)**. This collection of ligand fragments is clustered using a greedy clustering algorithm using RMSD to determine if two fragments are similar. If two fragments are within a 2.0 Å of each other, the fragment with a higher RMR6 is deleted. Remaining docked fragments are referred to as seeds **(e)**. The score distribution of a typical seed is given in **(f)** to show the exponential score shape of the distribution.

##### Dock and cluster rigid fragments

Intermolecular geometric and chemical complementarity between a protein and a ligand is essential for binding. Energetically preferred positions of ligand atom types can be captured using a discriminatory function (Figure 3c). Docking of seed fragments to the binding site grid is performed by moving seed’s rotational grid over the binding site grid points. Docked fragment poses that are in a steric clash with the protein are rejected (Figure 3d). A steric clash is considered if any interatomic distance between the fragment and the protein falls within nine-tenths of the atoms’ respective van der Waals sum. Each fragment translation and rotation that passes this initial filter is then evaluated with the RMR6 discriminatory function^56^. Finally, greedy clustering of docked and scored fragment poses in the Root Mean Square Deviation (RMSD) space computed based on their heavy atoms at 2 Å cluster cutoff is performed, resulting in a uniform distribution of locally best-scoring docked seed fragments covering the entire protein binding site (Figure 3e).

#### 2.2.3. Phase III: Flexible docking with iterative minimization

##### Generate partial compound conformations

For each compound to be docked, a user-specified percentage of each of its best-scoring rigidly docked seed fragment poses are considered. Among these, we search for such compatible pairs of docked seeds that are at the appropriate distances, that is, the distance between them is less than the maximum of their known bond distance. The maximum possible distance between a pair of seeds is calculated by traversing the path between the fragments in the original compound and summing up the distances between the endpoints of each rigid fragment on the path. We construct an undirected graph in which vertices represent seed fragments, and edges indicate that the corresponding pair of seed fragments is linkable. Using the MaxCliqueDyn algorithm^64^ we then find all fully connected subgraphs consisting of *k* vertices (k-cliques) in this graph, where the default value of *k* is set to three or to the number of seed fragments, whichever value is less. Each k-clique corresponds to a possible partial conformation of the docked seed fragments, in which these fragments are appropriately distanced so that they may be linked into the original compound. The possible partial conformations are then clustered using a greedy clustering algorithm at RMSD cutoff of 2 Å, where the best-scored cluster representatives are retained. The partial conformations sorted by their RMR6 scores from the best-to the worst-scored are used as an input to the next step of compound reconstruction.

##### Reconstruct compound with protein flexibility

Each identified partial conformation of the docked seed fragments is gradually grown into the original ligand by addition of non-seed fragments using the A* search algorithm. This can be done at different levels of protein flexibility. Protein minimization may be performed at each step of the linking process or only at the end when the compound has been reconstructed. Each seed fragment is linked to adjoining fragments according to the connectivity of the original compound. Each added non-seed fragment is rotated 360° about the bond vector at 60° increments. If the user has specified full protein flexibility, the resulting conformation of the partial compound and the protein is subjected to knowledge-based energy minimization using the RMC15 scoring function as for intermolecular forces. Simultaneously, bonds, angles and torsions of the partial compound and the protein are minimized using the standard AMBER molecular mechanics energy minimization. This procedure uses the popular OpenMM software package, specifically its implementation of the L-BFGS minimization algorithm^65^. With each round of minimization, the RMR6 score is calculated for the protein-compound interactions, and the scored conformation is added to the priority queue which consists of the growing compound conformations in the order from the best-scored to the worst-scored.

At each subsequent step of reconstruction, the A* search algorithm chooses the best-scored conformation from this priority queue and attempts to extend it. This conformation must meet an additional condition, which is that its attachment atoms that are to be connected by rotatable bonds to fragments not-yet added, need to be at appropriate distances from the attachment atoms on the remaining seed fragments. The algorithm iterates until the priority queue is empty in which case the compound has been completely reconstructed and is in a local minimum energy state. Alternatively, if the specified maximum number of steps was exceeded (1000 by default), then the reconstruction failed. The A* search is repeated for each partial conformation of docked seed fragments until all have been considered for reconstruction into a different docked conformation of the original compound. A final energy minimization procedure is performed on the protein-ligand complex treating the protein as fully flexible (side-chain and backbone) to remove steric clashes in the process of growing the ligand into the binding site. In addition to knowledge-based and molecular mechanics energy minimization, the fragment reconstruction process intrinsically accounts for ligand flexibility in the docking process. The described protocol results in a ranked list of docked and minimized protein-compound complexes.

### 2.3. Benchmarking the CANDOCK algorithm

#### 2.3.1. Benchmarking set of choice

We evaluated the CANDOCK hierarchical docking algorithm using a benchmarking set (1) to determine whether the algorithm can reproduce the crystal binding pose of the ligand in the binding site of the protein and (2) to correlate the scores of the three-dimensional (3D) docked poses of the ligand to the measured K_d_/K_i_ values of the ligand binding with the protein. The PDBbind benchmark^66,67^ is very well suited for this analysis because, for each protein in this set, it provides 3D coordinates and corresponding activity values for five protein-ligand complexes. In the CASF-2016 core set (previously referred to as PDBBind Core set v2016), there are a total of 285 such complexes for 57 proteins of interest to the medicinal chemistry community. The number of fragments present in a given ligand range from a single fragment to ligands consisting of thirteen fragments, enabling an evaluation of our method on both rigid and flexible ligands.

In addition to CASF-2016, we have also benchmarked our method against the Astex Diverse set^68^ as several protein-ligand complexes in this set include metal ions and other cofactors, allowing us to showcase these examples and assess how our algorithm handles these particular cases. We obtained each structure from the Astex set from the Protein Data Bank directly and only considered the biological assembly used to create the original benchmark.

#### 2.3.2. Input preparation

The binding site for both benchmarking sets is defined by spheres with a radius 4.5 Å centered around each atom of crystal ligand. We did not remove any cofactors, solvent molecules, ions, or glycans when preparing our docking runs. The provided reference ligand was used to generate fragments and seeds for docking.

#### 2.3.3. Parameters chosen for benchmarking

The most important parameter present in CANDOCK for linking seeds into ligands is the ‘Top Percent’ parameter as it is crucial to selecting the number of seeds used to generate potential conformations via the maximum clique algorithm^64^. If this number is too small, then there will not be enough potential conformations generated to sample the conformational space of the ligand properly. In fact, there is a possibility that no conformations are generated during the linking step, causing CANDOCK to fail to produce any conformations. If the ‘Top Percent’ is too large, then the conformational search space is too large, and CANDOCK will become computationally inefficient (especially in the case of fully-flexible protein docking). Therefore, we wanted to sample potential ‘Top Percent’ values to determine how well our method does at various levels of conformational space sampling. The values chosen for this parameter are 0.5%, 1.0%, 2.0%, 5.0%, 10%, 20%, 50%, and 100%. Default values of all the parameters used in the algorithm are listed in **Table S4**.

Similar to the conformational space sampled, we also investigated the effect of protein flexibility on the ability of the CANDOCK algorithm to reproduce the binding pose of a ligand. Accordingly, we used the algorithm in three modes: no protein flexibility (no energy minimization performed, maximum final iterations set to zero), with semi-flexible protein (final energy minimization only, default options), and with a fully flexible protein (iterative energy minimization performed, iterative flag turned on). The RMSDs for all poses generated from all ‘Top Percent’ values and all flexibility modes are calculated with respect to the experimental crystal pose using a symmetry independent method.

Finally, we determined the best scoring function to select the pose from all generated poses that best reproduces the crystal ligand pose (the ‘selector’ scoring function’) and potentially differentiate it from another scoring function used to rank the activity of a given ligand to the protein target of interest (the ‘ranker’ scoring function). To do this, we calculated the score of all poses generated for CASF-2016 using all scoring functions described in section 2.1. We then evaluated the ability of each scoring function to select the crystal pose of a ligand from all poses as well as the correlation between the score assigned to the selected pose and the experimental binding affinity. As there are 96 scoring functions, there are 9216 (96 ways to select by 96 ways to rank) different methods to rank the affinity of the ligands in CASF-2016. An overview of this benchmarking process for activity prediction is given in Figure 8.

## 3. Results and discussion

We discuss the performance of the CANDOCK algorithm in reproducing the crystal pose of a ligand via sampling the conformational space of the ligand in the binding pocket (including the entire chemical environment with cofactors, metal ions, crystal waters, etc.) modeled with different levels of protein flexibility for two benchmarking sets. In addition, we evaluate the ability of the algorithm to discriminate the crystal pose from all poses generated by the algorithm, and the ability to rank the activity of the ligands against the protein targets of interest.

### 3.1. Ligand conformational sampling is enhanced by fragment docking and protein flexibility

An important feature of any receptor-ligand docking methodology is its ability to generate docked crystal-like ligand poses within 2.0 Å RMSD of the experimentally determined pose of the native ligand. Using the CASF-2016 benchmarking set, we validated the ability of CANDOCK to generate crystal-like poses among the docked poses. We plotted the cumulative frequencies of all docked poses with the RMSDs from their corresponding crystal ligands’ poses for all ‘Top Percent’ values and for varying degrees of protein flexibility using the RMR6 scoring function (Figure 5; left-hand panels). Expectedly, these plots indicate that the use of larger (>20%) ‘Top Percent’ values generated significantly more poses within 2.0 Å than lower (<10%) ‘Top Percent’ values. For the semi-flexible (Figure 5c) method, the ‘Top Percent’ value of 20% yielded the highest number of poses within 2.0 Å of the crystal pose, with the corresponding cumulative frequency of ~91%, compared to independent benchmark of the best performing methods resulting in ~80% success to generate the pose^34^. The semi-flexible method thus outperformed the rigid protein (Figure 5a) and the fully flexible (Figure 5e) methods for the larger ‘Top Percent’ values that correlate with higher sampling of the ligand conformational space during fragment docking. However, the fully flexible protein method outperformed the semi-flexible (Figure 5c) and the rigid protein (Figure 5a) methods for smaller ‘Top Percent’ values such as 5% and 10%. In addition, the Boltzmann-like distributions in the RMSD plots (**Figure S1**) indicate that the CANDOCK algorithm adequately sampled the ligand conformations both far and close to the crystal ligand pose in CASF-2016. This suggests that the prediction of energetically-favorable ligand conformations is dependent on near-native protein flexibility during the linking of docked fragments. There are only 17 co-crystal structures (out of 285) where the semi-flexible algorithm failed to find a single crystal-like pose for the native ligand (1H22, 1H23, 1NVQ, 1U1B, 1YDT, 2P15, 2QNQ, 3AG9, 3BV9, 3KWA, 3O9I, 3PRS, 3UEU, 3URI, 3ZSO, 4EA2, 5C2H) for any top percent value. Additional 9 complexes (2C3I, 2CET, 2W66, 2WCA, 3ARU, 3BGZ, 3OZT, 3RR4, 3UEX) failed to find a crystal-like pose when the semi-flexible algorithm was used with a top percent value of 20%. Two of these complexes (3BV9, 3URI) contains a peptide ligand with a protein, a situation generally treated differently in other docking studies^34^. When fully-flexible docking is considered, CANDOCK fails on a total of 10 complexes, out of 285, resulting in an overall success rate of ~96% to generate crystal-like poses. Specifically, CANDOCK generates successful (crystal-like) poses for 7 complexes out of 17 failures from semi-flexible docking (3O9I, 2QNQ, 1YDT, 3ZSO, 5C2H, 3UEU, and 4EA2), and 2P15 becomes a near hit with an RMSD of 2.04Å. These results indicate that hierarchical generation of the ligand poses with the protein flexibility considered after fragment docking and ligand reconstruction is a successful strategy for enhanced sampling of the conformational space of ligands in protein-ligand complexes.

**Figure 4:**
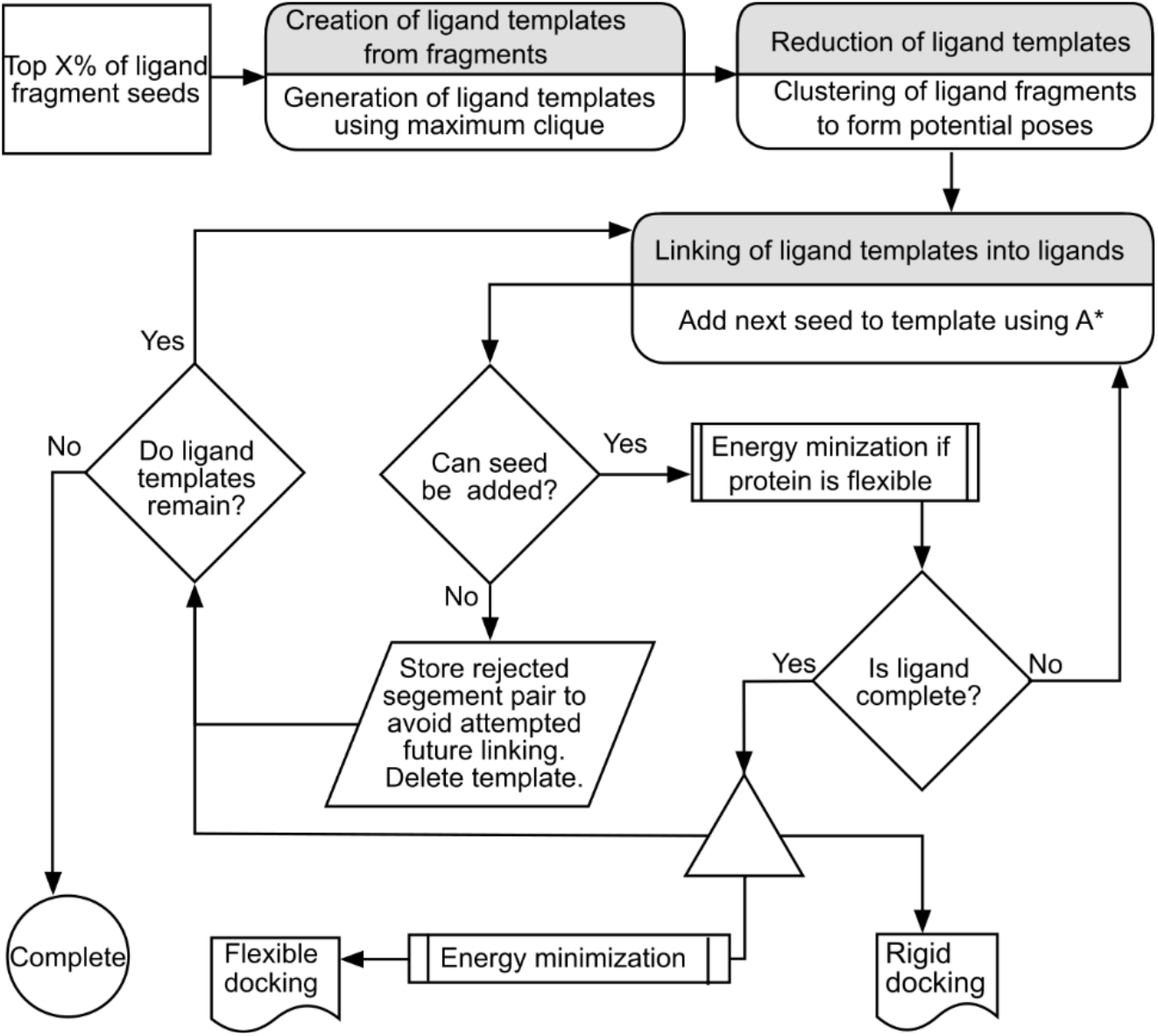
Workflow of the fragment linking procedure.

**Figure 5:**
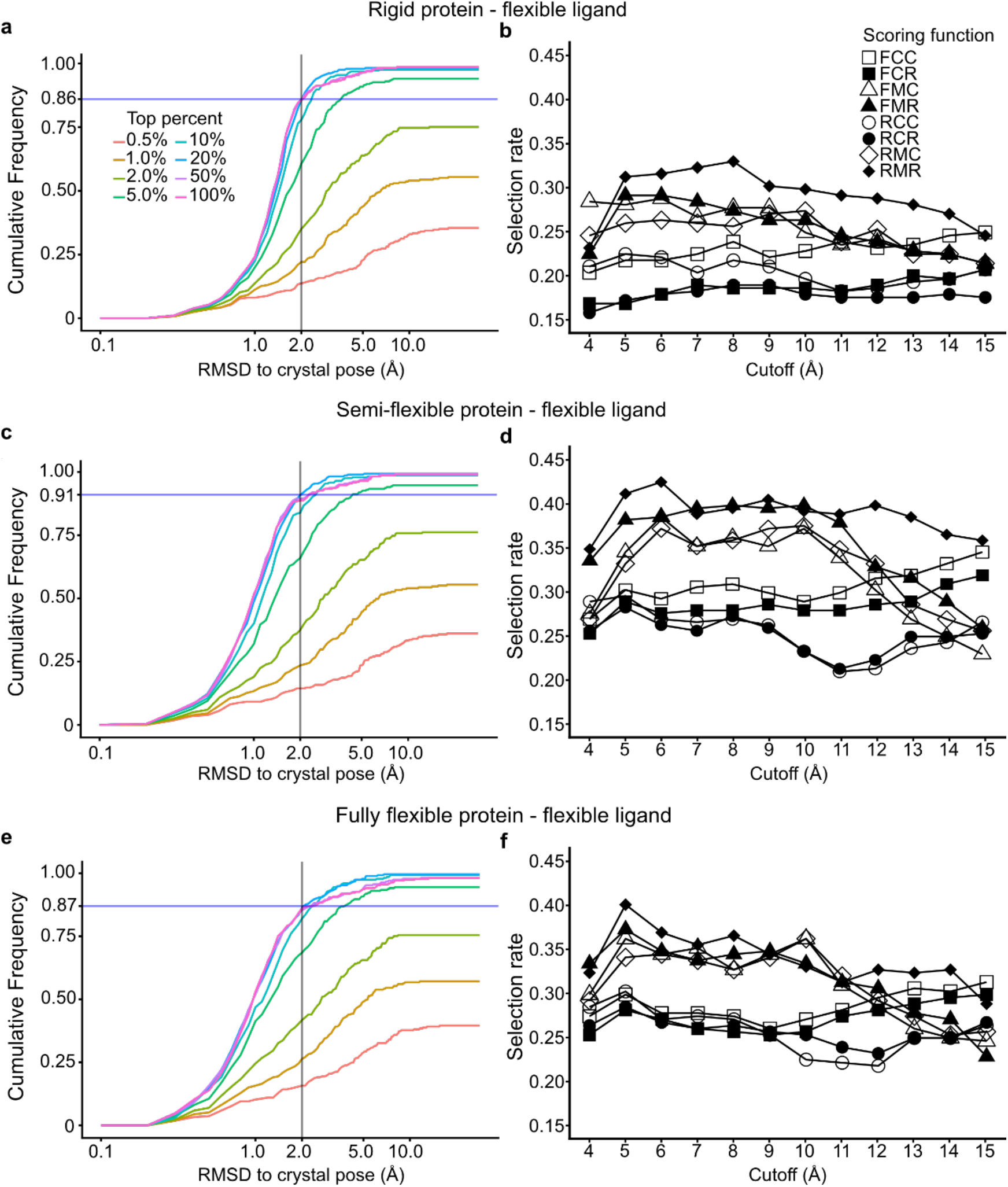
Cumulative frequency of the best RMSD pose generated by for rigid (flexible ligand only with no energy minimization of protein-ligand complex), semi-flexible (energy minimization of protein-ligand complex at the end), and fully-flexible (iterative energy minimization during linking procedure) CANDOCK docking results for the 285 proteins in CASF-2016 using the RMR6 scoring function are given in (**a)**, (**c)**, and (**e)** respectively. The selection rate, i.e., the portion of the best-scored docked poses within 2.0 Å of the crystal pose, is given for different scoring functions employed in (**b**), (**d**), and (**f**).

### 3.2. Radial Mean Reduced (RMR) scoring function family generates best docked ligand poses

We evaluated different scoring functions for their ability to select the crystal-like ligand pose as the highest-ranked pose, termed as ‘selectors’ henceforth. We calculated the selection rate for each scoring function at different radius cutoff values (Figure 5; right-hand panels) to identify best selectors. Here, the selection rate is defined as the fraction of the highest-ranked poses within 2.0 Å of the crystal ligand pose over all poses generated by the algorithm. The RMR family of scoring function at the cutoff radius value of 6 Å from each atom of the ligand, RMR6, performed best for the semi-flexible protein method, while the best selector scoring function for the rigid protein method was RMR8 and for the fully-flexible protein method was RMR5 scoring function. This shows that the RMR scoring function family is the best selector among 8 other generalized family of scoring functions. Conversely, the Radial Cumulative Complete (RCC) scoring function family performed the worst in selecting the crystal pose from the generated poses with the RCC11 scoring function being the overall worst selector. To elucidate the rationale behind the good performance of RMR6 in selecting a crystal-like pose, we plotted the RMR6 score of the docked ligands with lowest RMSD from the crystal pose against the RMR6 score of the crystal pose (**Figure S2**). For ‘Top Percent’ values >10%, there is a clear separation between the successful poses within 2.0 Å (blue points) and the failed poses far from the crystal ligand pose (red points). Moreover, these failed poses cluster above the diagonal line, indicating that RMR scores of failed complexes have higher energy value (as expected) than the crystal pose during sampling for ‘Top Percent’ values >10% (**Figure S2**). The number of failed poses decrease to lower numbers with increasing ‘Top Percent’, from 244 for 0.5%, 218 for 1.0%, 178 for 2.0%, 97 for 5.0%, 46 for 10%, 26 for 20%, 30 for 50%, and 32 for 100%. These data suggest a ‘Top Percent’ of 20% yields the highest number of poses within 2.0 Å of the crystal pose (previous section, Figure 5 - left-hand panels) and the number of failed cases are rare and clearly discriminated from both the crystal pose, as well as, the successful near-native docked poses (blue points) by using the RMR6 scores. Therefore, RMR6 can discriminate native and near-native interactions from a set of incorrect conformations generated by our docking method. Furthermore, RMR6 scoring function is a decent selector as the top pose (lowest RMR6 score) has an average selection rate of 41% for semi-flexible docking at a ‘Top Percent’ of 20% (Figure 5; right-hand center panels) and is comparable to the state-of-the-art independent benchmarks.^34^ Clearly, for some of the successful cases lowest RMR6 scores selected the pose within 2.0 Å RMSD of the crystal pose (**Figure S3**). However, RMR6 has a bias towards incorrectly scoring the top lowest scored RMR6 pose, better than the crystal pose for both successful and failed cases (blue and red points respectively are below the diagonal in **Figure S4**). If we include predicted poses other than the top pose, then we get a much higher selection success rate of 55% when top 2 poses are selected, 69% when top 5 poses are selected, and 76% when top 10 poses are selected. While the RMR6 scoring function is a decent selector, more work is needed to enhance the selection success rate, perhaps in combination with other scoring functions at different cut-offs along by using machine learning methods^69,70^. However, it is good to note that without any machine learning, our generalized RMR6 scoring function is comparable to successfully selecting a pose to a recently published neural network based scoring selection^71^ with a selection rate of ~50% for the top pose and ~65% for the top 5 poses. This suggests a reduced composition over all pairwise protein’s and compound’s specific atom types with mean reference state improves discriminatory accuracy by giving ‘context’ to the specific pose by solely including atom type interactions that are possible between the receptor and the ligand.

### 3.3. Docking long aliphatic chains needs enhanced sampling

We identified six complexes (1H22, 1H23, 3AG9, 3KWA, 3UEU, and 4EA2) out of 17 failed cases with CANDOCK semi-flexible algorithm with ligands that contain long aliphatic carbon chains (greater than 4 atoms). The remaining 11 complexes that fail are 3URI (8-mer peptide), 3O9I, 1U1B, 2QNQ, 3BV9 (6-mer peptide), 3PRS (14 fragments), 1YDT, 1NVQ, 2P15, 5C2H, and 3ZSO. If fully-flexible protein docking is considered, we get 4 complexes out of 10 failed cases that contain long aliphatic carbon chains (1H22, 1H23, 3AG9, 3KWA). CANDOCK does not consider aliphatic chain consisting of three carbon atoms (sp3 hybridized carbon; C3) as fragments for docking (see Materials and Methods). Instead, the A* search algorithm determines the docked positions by rotating them around the bond vectors of the growing chain at 60° increments. We hypothesize that this discrete sampling of conformational space, and not the potential functions in CANDOCK, is the cause for the poor performance of the algorithm on these compounds with many rotatable bonds. To test our hypothesis for the six failed long aliphatic carbon chain complexes (1H22, 1H23, 3AG9, 3KWA, 3UEU, and 4EA2), we scored the decoys provided by the CASF benchmarking set^67^ that included at least one pose within 2.0 Å RMSD. In all 6 cases, the RMR6 scoring function selected a pose within 2.0 Å RMSD of the crystal ligand, indicating that our generalized scoring function does not account for failure to identify crystal-like conformations (**Figure S5**). We plan to address this issue in detail in future versions of the algorithm by implementing a new sampling method or a ligand-class specific scoring function, similar to what was done for the support of carbohydrates in Autodock Vina separately^72^.

### 3.4. Full protein flexibility improves docking ligands with many rotatable bonds

The number of rotatable bonds in a ligand significantly influences the ability of docking algorithms to generate docked crystal-like ligand poses^34^. To study the effect of rotatable bonds on the performance of the algorithm, we compute the selection rate of the RMR6 scoring function against the number of fragments in a ligand (Figure 6). Due to the hierarchical fragment-based nature of the CANDOCK algorithm, the number of ligand fragments is used instead of number of rotatable bonds to measure CANDOCK’s performance. By comparing the fully-flexible protein method (Figure 6c) to the rigid protein method (Figure 6a) and to the semi-flexible method (Figure 6b), we show that the selection rate for flexible ligands increases with including protein flexibility during docking. Here, we define a flexible ligand with greater than 4 total fragments as the average number of fragments is 3.8 and the median is 3 fragments in the CASF-2016 dataset. Specifically, for the 216 ligands with four or fewer fragments, the semi-flexible (Figure 6b) and the fully-flexible (Figure 6c) methods performed equally well. The rigid, semi-flexible and fully flexible methods have a respective mean selection rates of 46%, 53%, 51% for the top pose; 61%, 65%, 65% when top 2 poses are selected; 74%, 77%, 81% when top 5 poses are selected; and 80%, 84%, 88% when top 10 poses are selected. Thus, full protein flexibility is not essential for ligands with less than 5 fragments as there is little difference in selection rate between semi-flexible and fully-flexible docking (Figure 6b,c). In contrast, for 69 ligands with greater than 4 fragments, the rigid, semi-flexible and fully flexible methods have a respective mean selection rates of 28%, 46%, 56% for the top pose; 35%, 59%, 68% when top 2 poses are selected; 44%, 75%, 84% when top 5 poses are selected; and 51%, 79%, 86% when top 10 poses are selected. Better performance of flexible methods versus the rigid method for larger ligands is most likely caused by the plateauing and even slight decline in the number of poses generated for ligands with >5 fragments for ‘Top Percent’ values >10% (**Figure S6)**. This suggests there is an upper limit to the sampling space possible for a given binding site and for a given ligand and once this limit is reached, the algorithm is no longer able to produce more docked ligand poses. However, the increased protein flexibility allows the CANDOCK algorithm to maneuver a larger ligand into a crystal-like binding pose, leading to higher selection rates observed for the semi and fully flexible protein methods.

**Figure 6:**
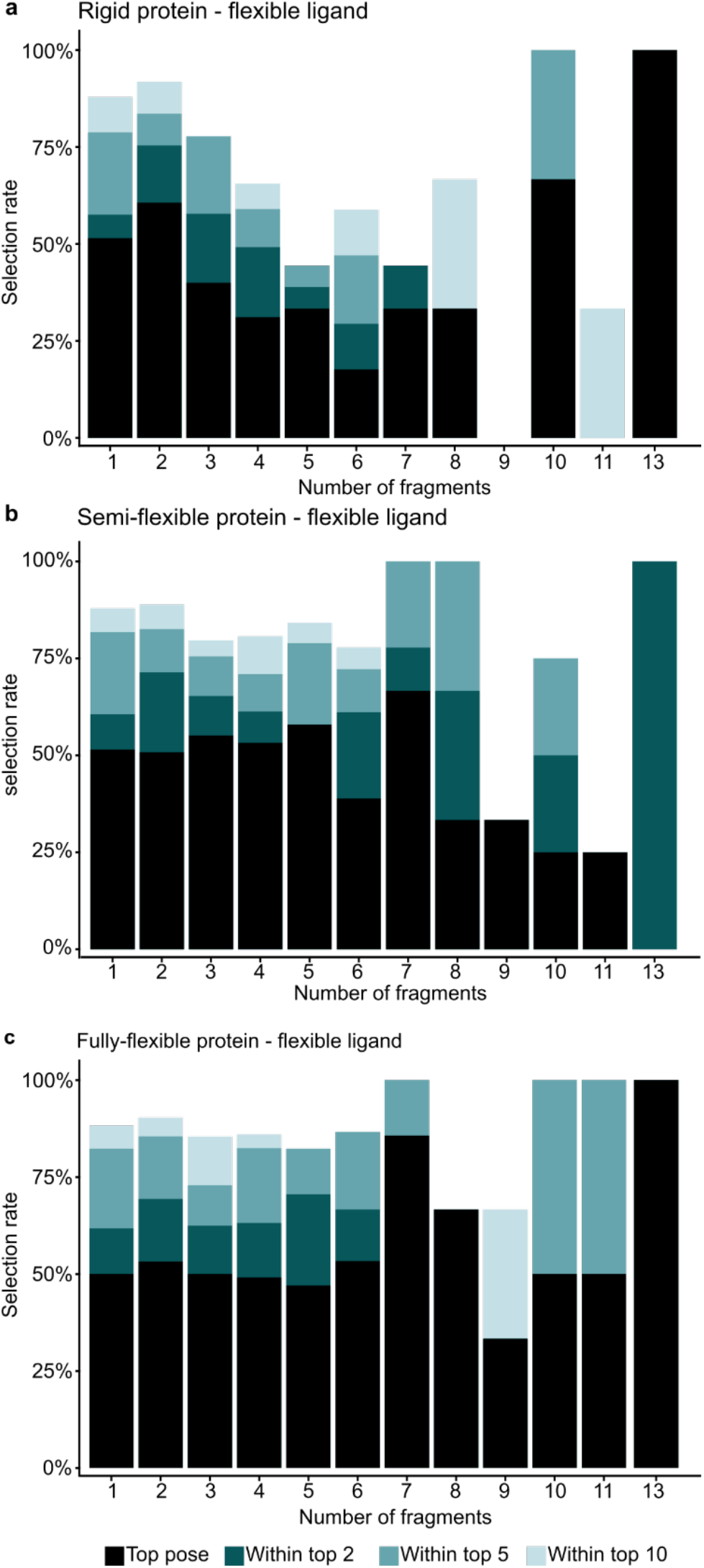
Selection rates for the RMR6 scoring function with rigid (**a**), semi-flexible (**b**), and fully-flexible (**c**) CANDOCK docking arranged by the numbers of ligand fragments in CASF-2016 (see Figure **2** for the definition of a fragment). For fragment counts greater than 13 (3URI, 3AG9, and 3PRS), CANDOCK did not produce any poses within 2.0 of the crystal pose.

### 3.5. Inclusion of chemical environment and cofactor interaction in binding sites lead to accurate crystal-like ligand pose generation

The Astex Diverse Set^68^ is a widely used benchmarking set for measuring a docking program’s ability to predict the native pose of a ligand. One important feature of this set, compared to CASF-2016^67^, is the inclusion of several cofactors and metal ions such as zinc ions and heme groups in the binding sites. Traditionally, with docking methods, the cofactors in the binding pockets have been ignored or treated as non-physical models with improper representations that affected performance^67^. As an example, for Heme groups, we used a previously published extension to the GAFF forcefield to ensure proper representation of this cofactor during the minimization procedure^73^, compared to other methods treating it as a hydrogen bond donor^24^. We hypothesize that in order to perform well on this benchmarking set, the docking algorithm must properly sample ligand conformations interacting with metal ions and doing so requires adequate representation of metal-ligand interaction potentials at the atomic scale. A generalized potential function can include all relevant cofactors, metal ions, etc. in the binding pocket as separate interactions (**Figure S7**), compared to one metal-ion type used by others^24,67^. To highlight the ability of our scoring function to characterize such interactions in a pair-wise fashion, we plotted various atom pair interactions of interest to medicinal chemists (**Figure S7**).

The number of complexes in this benchmarking set where CANDOCK algorithm produces a ligand pose within 2.0 Å RMSD of the crystal pose is given in Table 1. CANDOCK successfully generates a crystal pose for 97.6% of the Astex benchmarking set (83 out of the 85 complexes). We attribute this success to the ability of our algorithm to properly sample the conformational space of ligand in the binding pocket while considering all interactions of the ligand within the binding pocket including cofactors, metal ions, etc. In a recent comparison using Astex dataset^28^, the success rate for FlexAID^28^, Autodock Vina^24^, FlexX^78^, and rDock^47^ are 66.7%, 81.8%, 78.8%, and 89.4% respectively, when all 85 complexes are considered. When 16 complexes containing a metal ion were removed (1GKC, 1HP0, 1HQ2, 1HWW, 1JD0, 1JJE, 1LRH, 1MZC, 1OQ5, 1R1H, 1R55, 1R58, 1UML, 1XM6, 1XOQ, 1 YQY), the success rates of these methods increased to 72.1%, 83.6%, 79.7%, and 91.3% respectively^28^. CANDOCK outperforms these methods without removing metal ions complexes from the benchmarking set, supporting the hypothesis of adequate sampling and included proper representation of interactions within the binding site. The two complexes where CANDOCK nearly missed to generate a crystal pose using the semi-flexible method are 1HP0 (lowest RMSD of 2.08) and 1W1P (lowest RMSD of 2.734). Additionally, when the protein is considered as a rigid body (rigid docking), CANDOCK failed to find crystal poses for 1Y6B and 1MZC as well (81 out of 85 complexes in Table 1). The algorithm also performs well on complexes that failed by using other popular docking methodologies for the Astex Diverse set. According to a previous study^28^, there are four complexes (1G9V, 1GM8, 1JD0, and 1MEH) where Autodock Vina^24^, rDock^47^, FlexX^78^, and FlexAID^28^ all have difficulty reproducing the crystal-like pose of the ligand but CANDOCK successfully generated a crystal-like pose. The interactions of the ligand with cofactors in the binding pocket for these complexes are shown in **Figure S8**. Specifically, 1G9V have cation-**π** interaction and 1GM8 have **π**-**π** interactions between an aromatic ring and the surrounding protein environment. Similarly, 1MEH contains a **π**-**π** stacking interaction between the ligand and a cofactor. 1JD0 has an interaction between the zinc ion and a sulfonyl group. These complexes showcase the success of our hierarchical docking method over previously published works.

**Table 1.** Number of successes in the Astex diverse Set for all ‘Top Percent’ values investigated. There is a total of 85 protein-ligand complexes in this benchmarking set.

| **Top percent** | **Rigid Protein** | **Semi-Flexible Protein** |
| --- | --- | --- |
| **0.5%** | 7 | 7 |
| **1.0%** | 14 | 15 |
| **2.0%** | 28 | 33 |
| **5.0%** | 57 | 60 |
| **10%** | 67 | 74 |
| **20%** | 77 | 79 |
| **50%** | 79 | 82 |
| **100%** | 78 | 81 |
| **ALL POSES** | 81 | 83 |

We also consider specific cases where cofactors interaction with the ligand in a given complex successfully reproduced the crystal pose (Figure 7). Specifically, in Figure 7a-b, for oxygen-zinc interactions in 1HWW and 1R55 during docking, the energy minimization procedure moved the location of the Zn^2+^ ion in the binding pocket (2.4 Å and 1.5 Å respectively) as there are no constraints to restrict its movement within the binding pocket. This movement does not prevent the algorithm from generating a pose within 2.0 Å RMSD of the native structure. For 1OQ5 and 1JD0, the docked poses of ligands interacts with a zinc ion through a sulfonyl amide group (Figure 7c-d) and it is interesting to note that the zinc ion moved much less in these cases (0.5 Å and 0.6 Å). For the ligand in 1OQ5 (Figure 7c), the orientation of the sulfonyl amide aligns perfectly with the reference crystal pose, suggesting that the interactions with sulfonyl amide group caused the zinc ion to stay in place. For the ligand in 1JD0 (Figure 7d), the docked pose of the same group does not align with its reference; however, the overall pose still is within 2.0 Å of this reference. Therefore, the ability for the algorithm to produce a pose within 2.0 Å of the reference is not dependent on correctly predicting the orientation of all functional groups in a given molecule.

**Figure 7:**
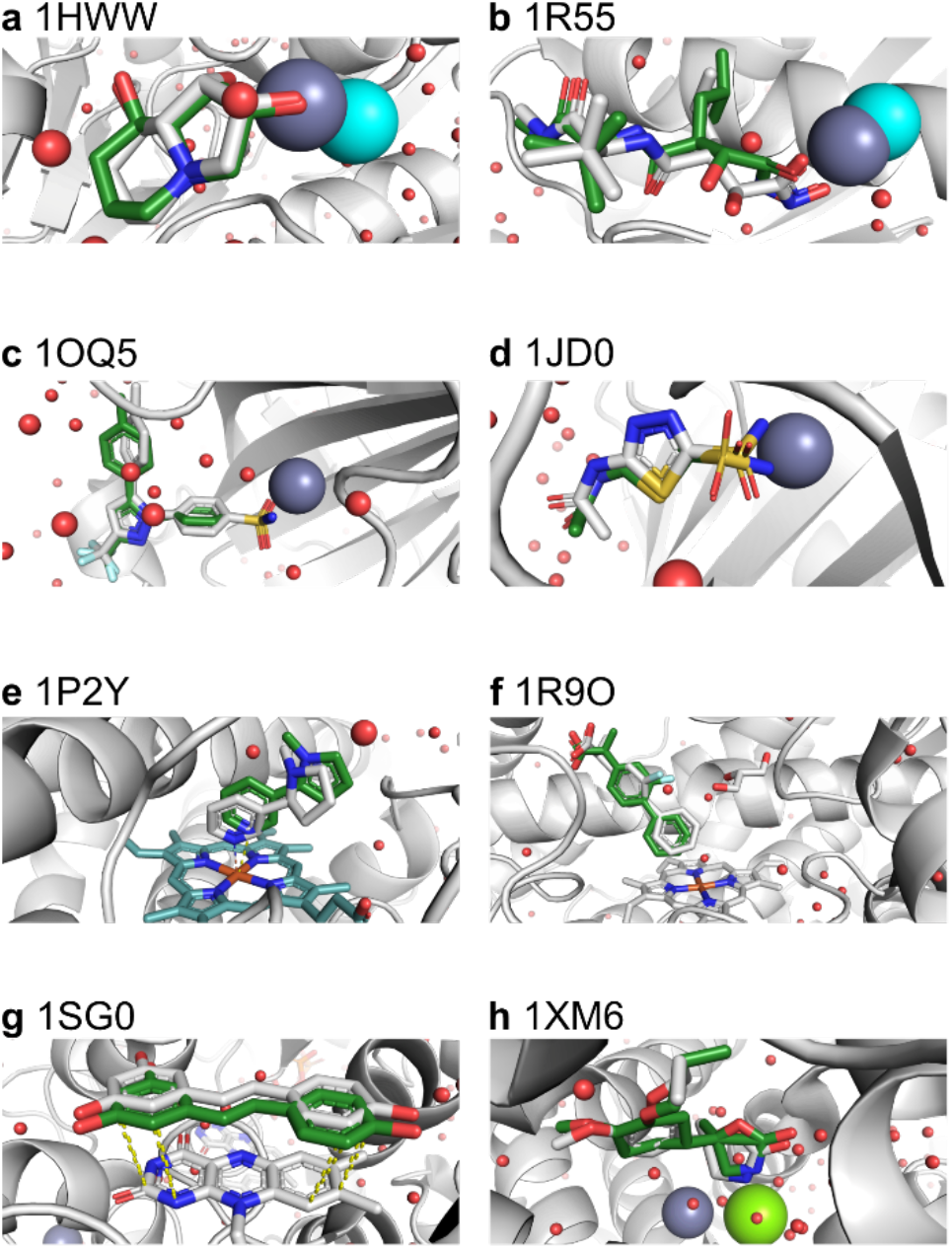
Example docked ligand poses from the Astex Diverse set that show versatility of the CANDOCK algorithm in handling cofactors. In all panels, the reference pose is given in white and the lowest RMSD pose predicted by CANDOCK with a ‘Top Percent’ value of 20% using the semi-flexible method is given in green. Panels (**a**) and (**b**) were selected due to presence of oxygen-zinc interactions between native ligand and protein. The zinc ion before and after energy minimization is given in gray and cyan respectively showing that the energy minimization moved the zinc ion considerably. The complexes in (**c**) and (**d**) show the interactions between sulfonyl amide groups and a zinc ion. The interactions of a compound with a heme group via a nitrogen lone pair is shown in (**e**) and the interaction of an aromatic carbon with a heme group is given in (**f**). Finally, panels (**g**) and (**h**) show the interactions of compounds with other cofactors, such as a π-π interaction of a compound with flavin-adenine dinucleotide and interaction of a compound with zinc and magnesium in a binding pocket.

We selected a larger organic cofactor (heme group) in the binding site of the protein-ligand complexes, 1P2Y and 1R9O (Figure 7e-h). The heme group is present in several liver enzymes^74–76^, therefore predicting the location of a ligand relative to this group is important for medicinal chemistry. For 1P2Y, CANDOCK predicts the pose of a compound relative to the heme group when the nitrogen of the compound is interacting with the iron atom of this group (Figure 7e). Similarly, for 1R9O, a successful pose is generated including the interaction between an aromatic carbon and the iron atom (Figure 7f) indicating that proper representation of heme group is essential to capture such interactions to generate the binding pose. We also demonstrate that generating a crystal-like docked ligand pose in the presence of a large cofactor is independent of the size of the cofactor itself. This is shown for 1SG0 complex containing the flavin-adenine dinucleotide cofactor (Figure 7g) where the dominant interaction between the ligand and the cofactor is π-π stacking. A crystal-like pose was also reproduced when the type of interaction changed dramatically, as shown in 1XM6 for the binuclear metal center formed by zinc and magnesium ions (Figure 7h). These interactions are important for developing phosphodiesterase inhibitors^77^, therefore it is encouraging to observe CANDOCK’s ability to reproduce a crystal pose in these cases. We conclude that the algorithm is able to generate a crystal-like docking pose by including interactions with diverse cofactors in the binding pocket.

### 3.6. Radial Mean Complete (RMC) scoring function at 15 Å cutoff is best for energy minimization

A potential or scoring function, used for energy minimization of a protein and a ligand should correlate quantitatively with the RMSD between the docked ligand and the crystal ligand, so that a decrease in score corresponds to a decrease in RMSD. Therefore, to determine the best minimization function, we calculated these correlations expressed as the average and the median Pearson correlation coefficients for all the scoring functions evaluated over CASF-2016 (**Table S1**). **Figure S9** shows that the RMC and FMC scoring function families have the largest correlation with RMSD (average across all cutoffs is 0.30 units greater than averages for other scoring functions). Moreover, with increase in the cutoff value for RMC and FMC scoring functions, the correlation also increased from an average of 0.36 at 4 Å to an average of 0.56 at 15 Å suggesting that including long-range interactions is essential. We also show that that the median and the average of these correlation values for the RMC and FMC scoring function families are relatively similar, indicating that the distribution of correlation values is not biased towards high or low correlations for any given protein in the CASF-2016 set. In addition, the RMC15 score of the experimental crystal pose has a strong correlation with the RMC15 score of the lowest RMSD pose (**Figure S10**, r^2^ > 0.99). Finally, the pose with the lowest RMC15 score correlates well with the RMC15 score of the crystal pose (**Figure S11**, r^2^ > 0.95). Taken together, we conclude that using the RMC15 scoring function in the CANDOCK algorithm to calculate intermolecular forces and energies during the energy minimization of the docked protein-ligand complexes correlates well with RMSD from crystal ligand pose (few example cases of RMSD vs RMC15 score plots are shown in **Figure S12**).

### 3.7. Crystal pose prediction method is independent of ranking ligand binding affinities

Another critical aspect of the scoring function is the ability to accurately rank the relative binding affinities of known binders to the same protein target. A stringent criterion for testing the ranking ability of a scoring function is by docking the compounds to the targets and compare to experimental binding affinities, i.e. without knowing the crystal pose of the ligand. CASF-2016 provides experimental binding affinities (pKi/pKd) and three-dimensional coordinates of 57 protein targets with 5 compounds each for a total of 285 pKi/pKd values for protein-ligand complexes. We determined the correlation between the 285 experimental binding affinities (pKi/pKd) with docking scores for 285 docked poses selected using each of the generalized scoring functions. We found that RMR6, our best ‘selector’ scoring function for selecting the crystal-like pose, does not adequately correlate with the pKi/pKd values supplied by CASF-2016 (Pearson correlation of −0.38) suggesting a need for a different scoring function for scoring the crystal-like selected pose. Therefore, we developed a procedure (Figure 8) to first select the representative docked pose of a complex using a scoring function (selector) and then rank using another scoring function (ranker) to obtain a good correlation with the pKi/pKd values.

**Figure 8:**
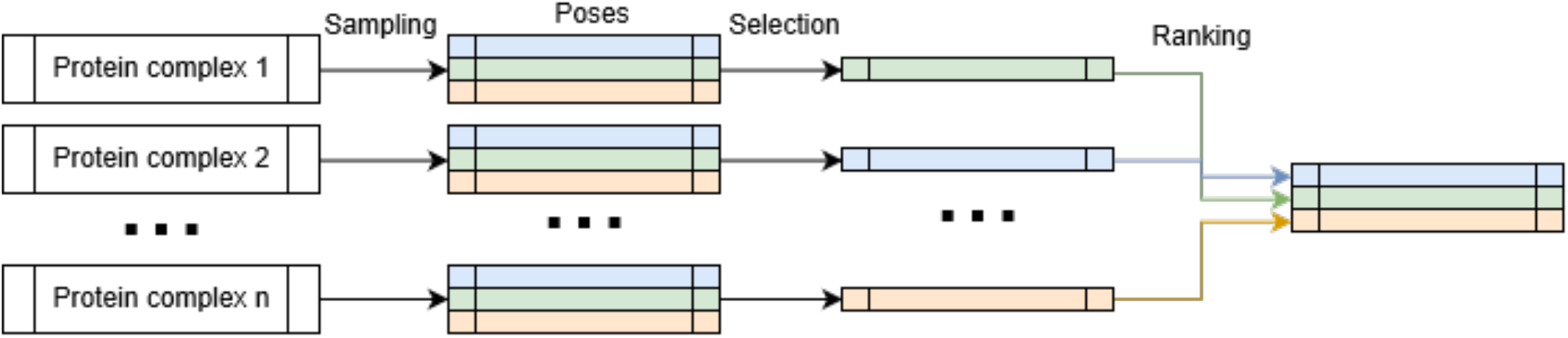
CANDOCK activity evaluation pipeline. Sampling is performed using the RMR6 scoring function to generate thousands of ligand poses. The best pose is selected with a ‘selector’ scoring function to represent the protein-ligand complex. Only this selected pose is rescored using the ‘ranker’ scoring function, which is used to assign a new score to the complex. The best ranker score on the selected pose is used to rank the protein-ligand complex based on correlation with pKd/pKi data.

The best ‘ranker’ scoring functions are RMC15 and FMC15 (Figure 9a and 9b) that were selected based on both Pearson and Spearman correlation between all 96×96 selector and ranker scoring functions combinations with the experimental pKi/pKd data in CASF-2016. There was little difference between the worst crystal pose selector (RCC11 that selects top pose 22% of the time, Figure 9c) and the best selector (RMR6 that selects top pose 43% of the time, Figure 9d), indicating that the ability of a selector to find the crystal-like pose is not important for correctly ranking the binding affinity of the ligand. This is also evident as the difference in correlation for the worst (RCC11) and the best (RMR6) selectors in combination with the best ranker (RMC15) score is 0.024. Furthermore, the correlation between the RMC15 score (best ranker) and the pKi/pKd data for all 96 possible selectors (shown in Figure 9e) have a small deviation (standard deviation of 0.0829 for the average Pearson correlation). This suggests that the selection of the pose has a minor impact on ranking the activity of the ligand. This result is further supported by **Figures S13-S17** and **Tables S2-S3** where the selector is either the best-scored pose using RMR6 scoring function or the lowest RMSD pose from the crystal ligand. We find that either of these selectors do not improve the ability of the best ranker (RMC15) scoring function to rank the pKi/pKd data of compounds binding to the same protein. Additionally, the difference in the overall Pearson correlation for the minimum RMSD pose selector vs the RMR6 pose selector is 0.001. Finally, it is important to note that the RMC15 score of weak binders in CASF-2016 (pKi < 2.5) does not correlate similar to the remainder of the poses (Figure 9c-d) as removal of these ligands increases the correlation between the RMC15 score and binding affinity by 0.241. While these findings are encouraging as they suggest to remove the burden of finding the crystal pose of the ligand, a more detailed study with an additional benchmarking set, such as the Directory of Useful Decoys (DUD-E)^79^, is required to determine the proper choice of scoring function or combinations to rank protein-ligand complexes and discriminate weak and non-binders will be addressed in future work.

**Figure 9:**
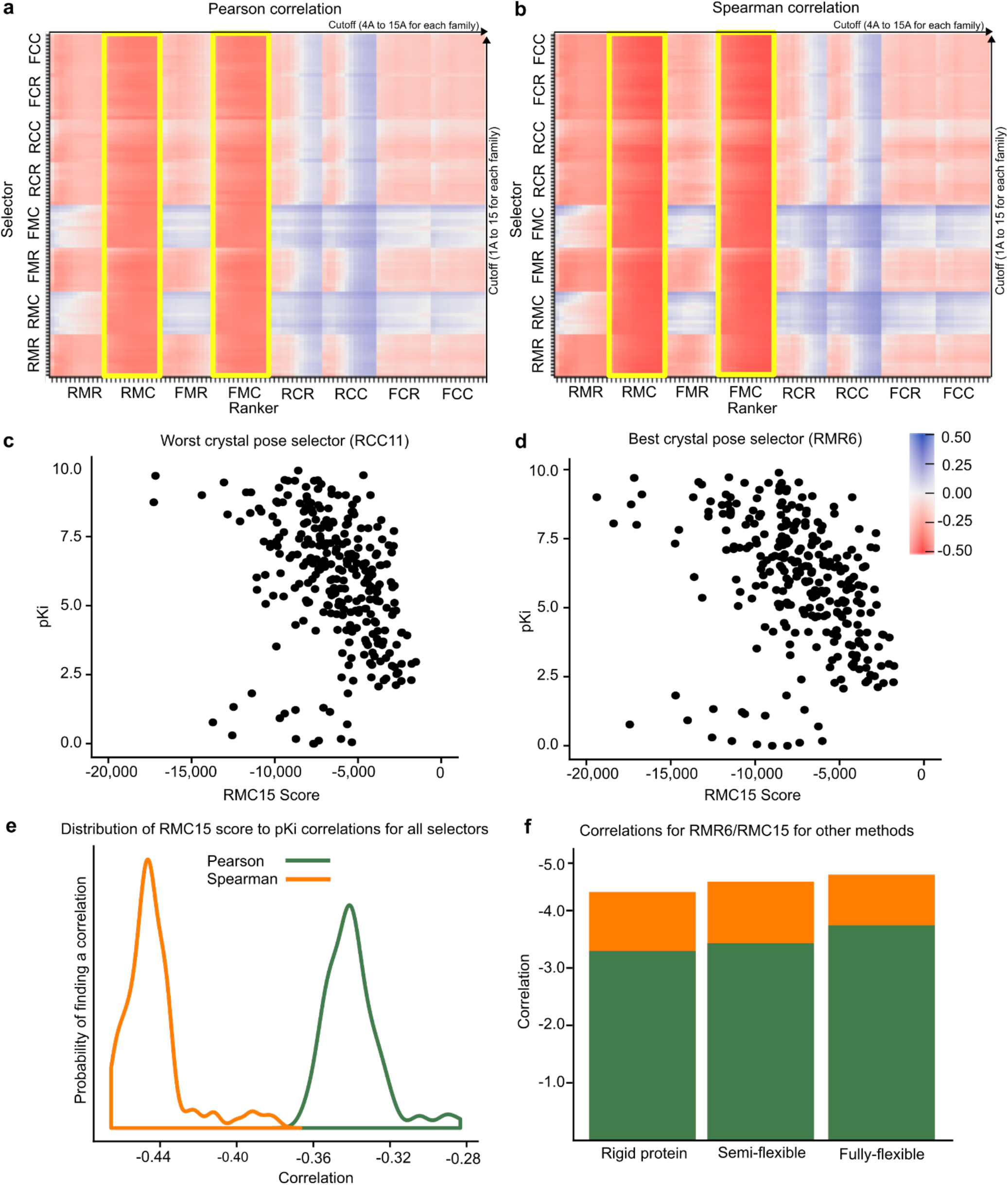
The Pearson (**a**) and Spearman (**b**) correlation coefficients between all pairs of selector and ranker scoring functions (arranged by family) and the experimental pKi of any complexes in CASF-2016. Note a negative correlation between score and pKi/pKd is expected as the ‘p’ operator introduces a negative sign to the affinity (the smaller the Ki, the larger the pKi). The RMC and FMC (highlighted in yellow) families perform best and there is a general trend where an increase in cutoff (from left to right) results in improved performance in ranking complexes in order of their measured pKi. Plots of pKi vs. RMC15 score are given in (**c**) and (**d**) for the worst crystal pose selector (RCC11) and the best crystal pose selector (RMR6), respectively. The lack of major differences between these two selectors with the same ranker indicates the lack of importance in selecting the correct binding pose for ranking the pKi of a protein-ligand complex. (**e**) The distribution of all correlations, regardless of selector, for the RMC15 scoring function (**f**) The correlations for other docking methods with RMR6 as the selector and RMC15 as the ranker.

Similar to the selector used, the flexibility mode (rigid, semi-flexible, fully-flexible) used to generate ligand poses does not have a significant impact on the correlation between score and binding affinity (see Figure 9f). While the fully-flexible methodology has a significant advantage for the kinases such as, ABL1, JAK2, and CHK1 (**Figure S15**), there are many other examples of protein-ligand complexes where the semi-flexible method provides a clear advantage over the fully-flexible and rigid methodologies (**Figures S13-S14**). This is significant because semi-flexible method is less computationally demanding than the fully-flexible method and can be used efficiently in a virtual screening pipeline. Moreover, in some cases, the correlations between the scores and pKi/pKd data have variability based on the type of protein. For example, the nuclear hormone receptors ER and AR have positive correlation values instead of the expected negative ones; the best selector/ranker pair for HIV proteases in CASF-2016 is RMC15/RMR6 which is the opposite of what was found for other test cases of CASF-2016, in general. Therefore, the use of different scoring functions for different protein classes may be advantageous in ranking the relative binding affinity of the ligands to the protein targets, which remains to be studied in our future work.

## 4. Conclusions

We present the CANDOCK algorithm, our hierarchal atomic network-based docking algorithm that accounts for protein flexibility and ligand interactions with all cofactors, metal ions, etc. in the binding pocket using generalized statistical scoring functions. We demonstrated that these scoring functions worked very well to generate a crystal-like pose for ~94% of the CASF-2016 dataset consisting of 285 protein-ligand complexes. There were 17 (of 285) failures in total with semi-flexible docking, which were reduced to 10 failures with fully flexible including 4 (out of 10) failures that contain long aliphatic chains. We found that RMR6 scoring function was the best at selecting a crystal-like ligand pose and RMC15 scoring function scored the selected poses to rank ligands according to their measured binding affinities. Our algorithm only requires a final energy minimization of the protein and the ligand (semi-flexible) to generate crystal-like ligand poses for ligands consisting of less than six fragments, compared to fully-flexible methods needed for larger ligands. CANDOCK was developed to provide proper representations of ligand, receptor, and all cofactors in the binding pocket. It performs well by including ligand and cofactors interactions in the binding pocket using the generalized statistical potential and without the need for parameterization. CANDOCK successfully generates a crystal pose for 97.6% of the Astex benchmarking set (83 out of the 85 complexes) that includes generating crystal-like poses for cases that failed with all popular docking methods (e.g. containing metal-organic interactions). We show that the RMR6 scoring function using a short distance cutoff and reduced atom type set is adequate for selecting the crystal pose of the ligand. However, a longer distance cutoff and complete atom type set used in the RMC15 scoring function are essential to achieve reasonable correlation between the docking score and the RMSD of a docked ligand from the crystal ligand, which justifies the use of RMC15 as the minimization function. The RMC15 scoring function was also the best at reproducing reasonable correlations between scores and ligand binding affinities. We believe that the release of the CANDOCK algorithm will give the community a valuable freely available tool for generating chemically relevant ligand poses for use in drug discovery efforts. The hierarchical nature of our method presents a powerful and flexible tool to performs proteome-wide docking studies efficiently, yielding an improved drug discovery and design pipelines.

