## Supporting Information for "CANDOCK: Chemical atomic network based hierarchical flexible docking algorithm using generalized statistical potentials"

[8]Integrative Data Science Initiative

[‡]These authors share equal contribution to this work.

*Corresponding Author

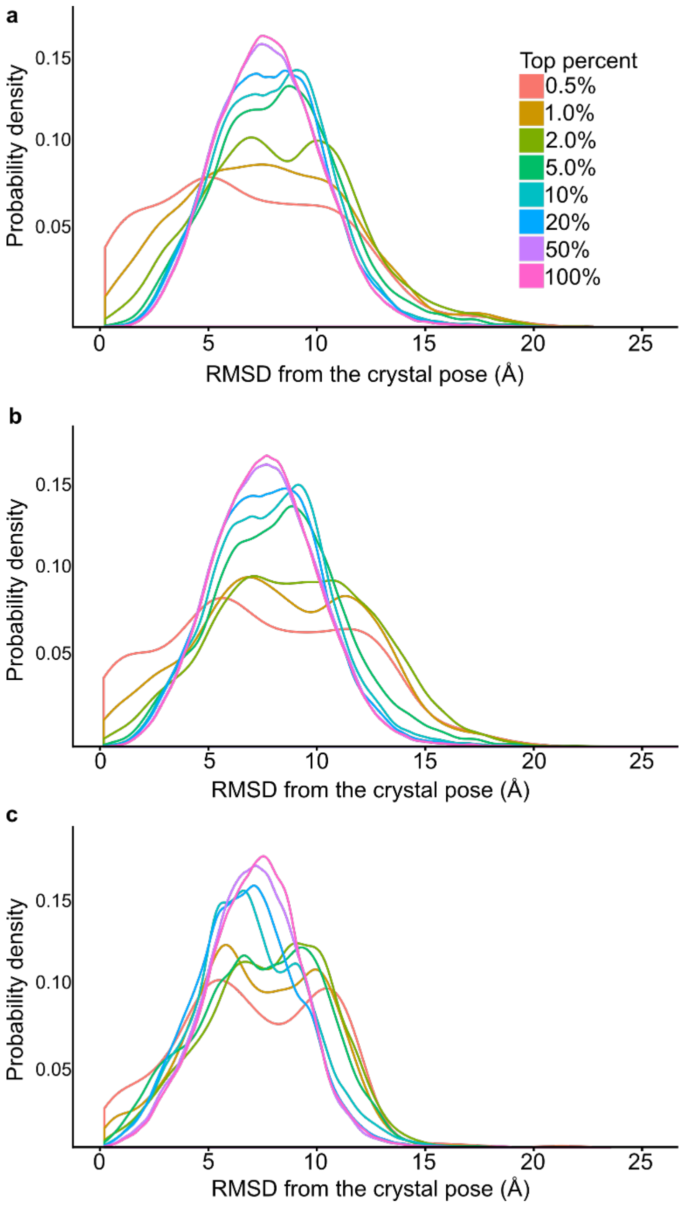


**Figure S1:** Distribution of RMSD values (Å) for all ligand poses generated by CANDOCK for docked poses in the CASF-2016 benchmark for **(a)** rigid-protein docking, **(b)** semi-flexible protein, and **(c)** fully-flexible protein docking.

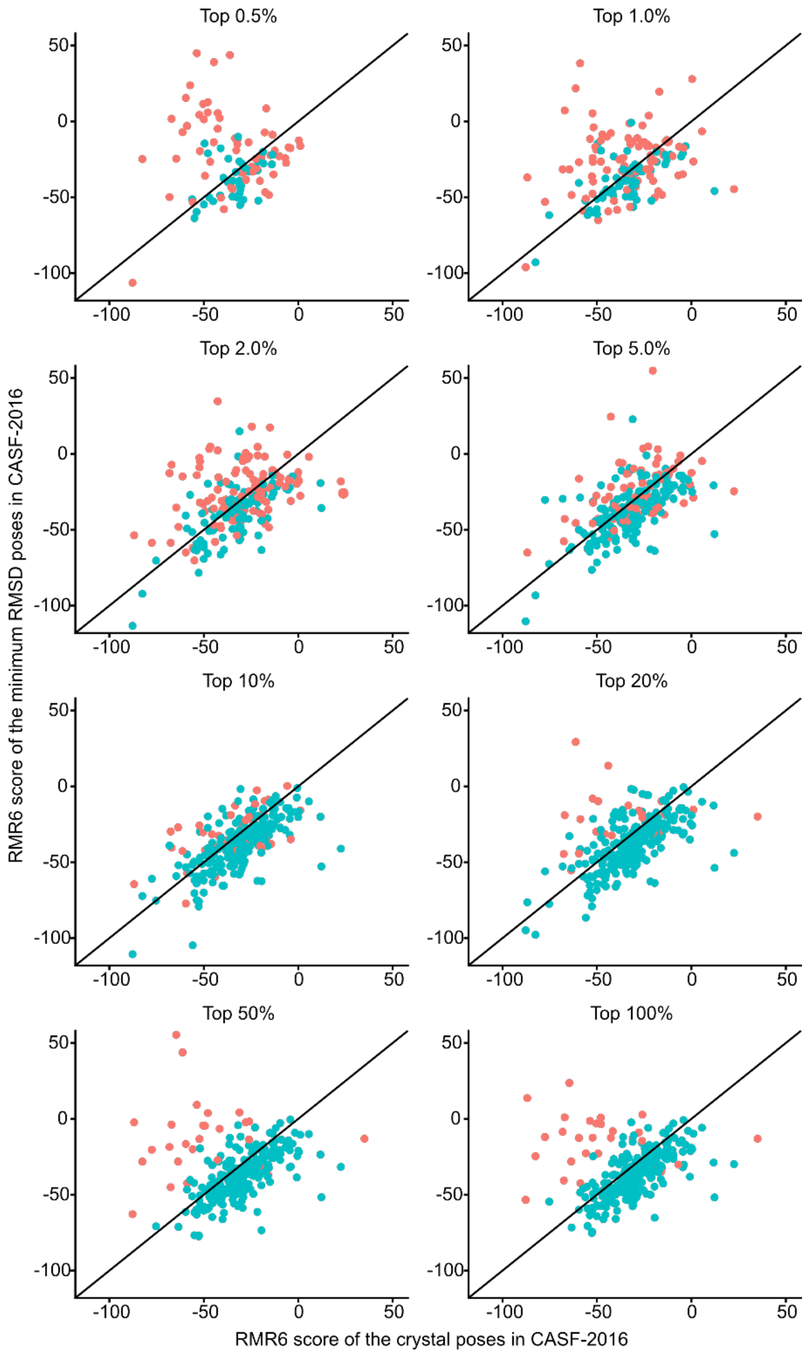


**Figure S2:** Correlations between the RMR6 scores of the crystal poses and the pose with the lowest RMSD are shown for all eight top percent values for complexes in CASF-2016. Poses within 2.0 Å of the crystal pose are shown in blue (success) while poses with RMSD >2.0 Å (failures) are shown in red. For top percent values greater than 20%, the complexes that failed cluster above the y=x line. Therefore, in these cases, the CANDOCK algorithm did not sample the conformation space close to the binding pocket.

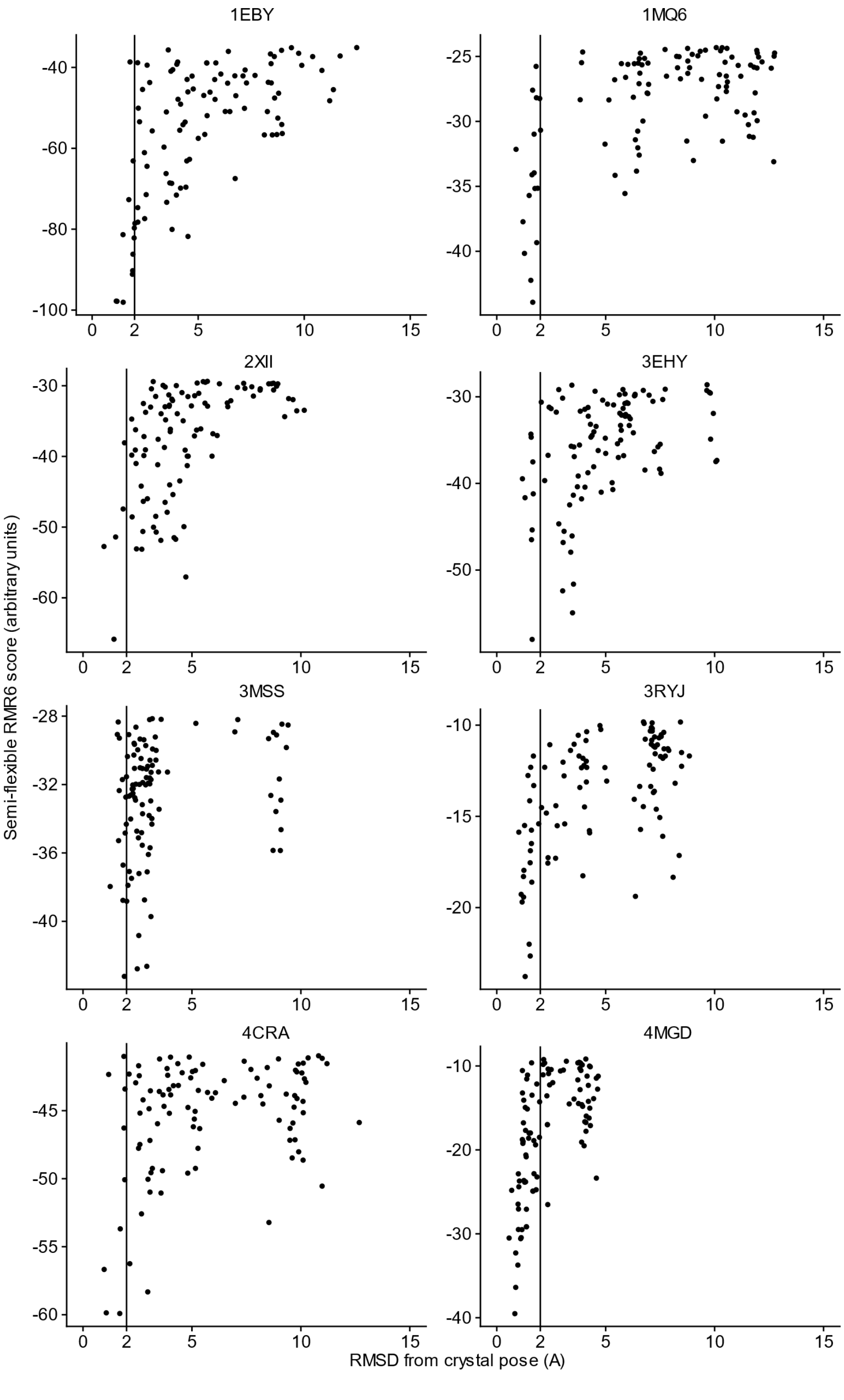


**Figure S3:** Plots of the RMR6 score of all poses produced by CANDOCK for selected proteins in CASF-2016 versus the RMSD of the pose. In all plots, the RMSD ranges from 1Å to 15Å. The poses were obtained using the semi flexible method at a 'Top Percent' value equal to 20%. These of these plots show a tunnel like affect around as one approaches an RMSD of zero, showing the scoring functions ability to select the crystal pose in these cases.

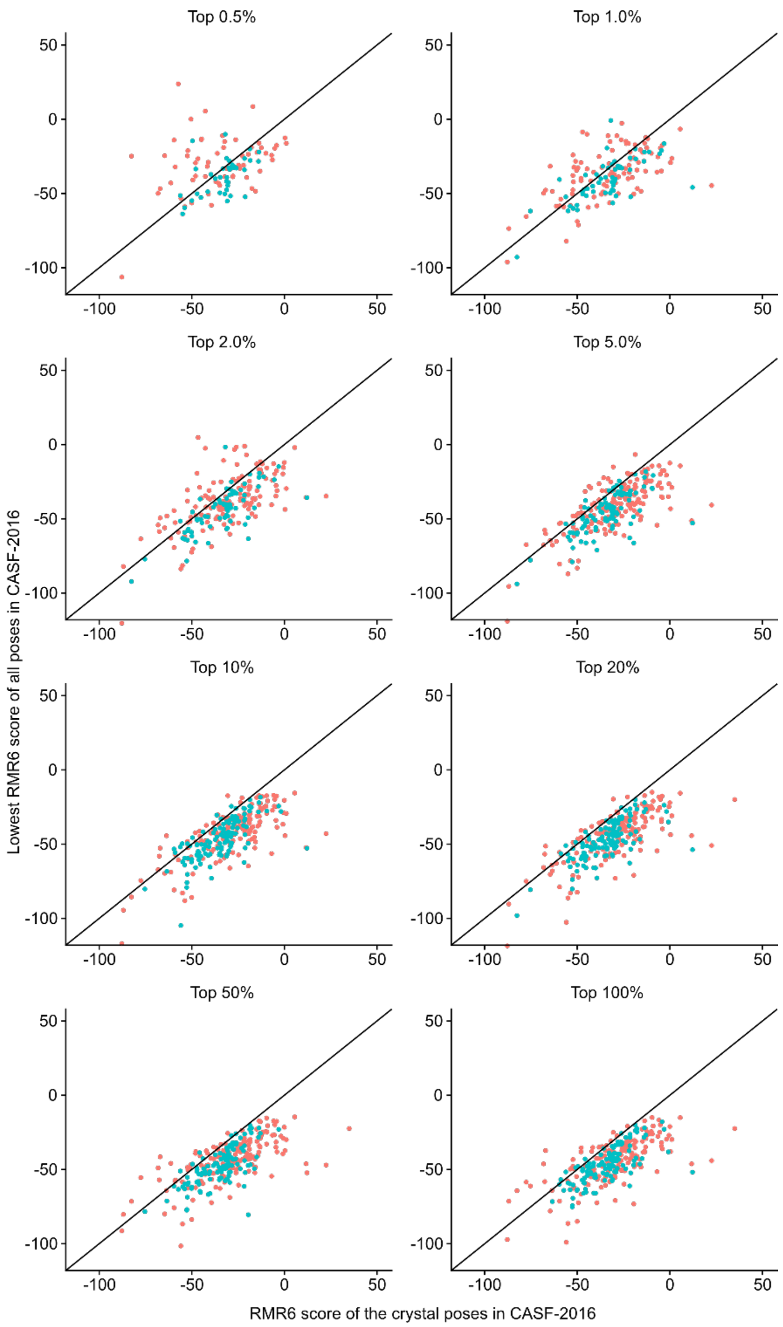


**Figure S4:** The lowest RMR6 score obtained for each cocrystal is plotted against the RMR6 score of the crystal pose. Poses within 2.0 Å of the crystal pose are shown in blue (success) while poses with RMSD>2.0 Å are shown in red. The majority of points on this graph cluster below the y=x line, indicating that the RMR6 scoring function incorrectly scores several poses more favorably than the crystal pose, regardless of if the pose is close to the crystal pose. Therefore, there are potential improvements to be made for this scoring function.

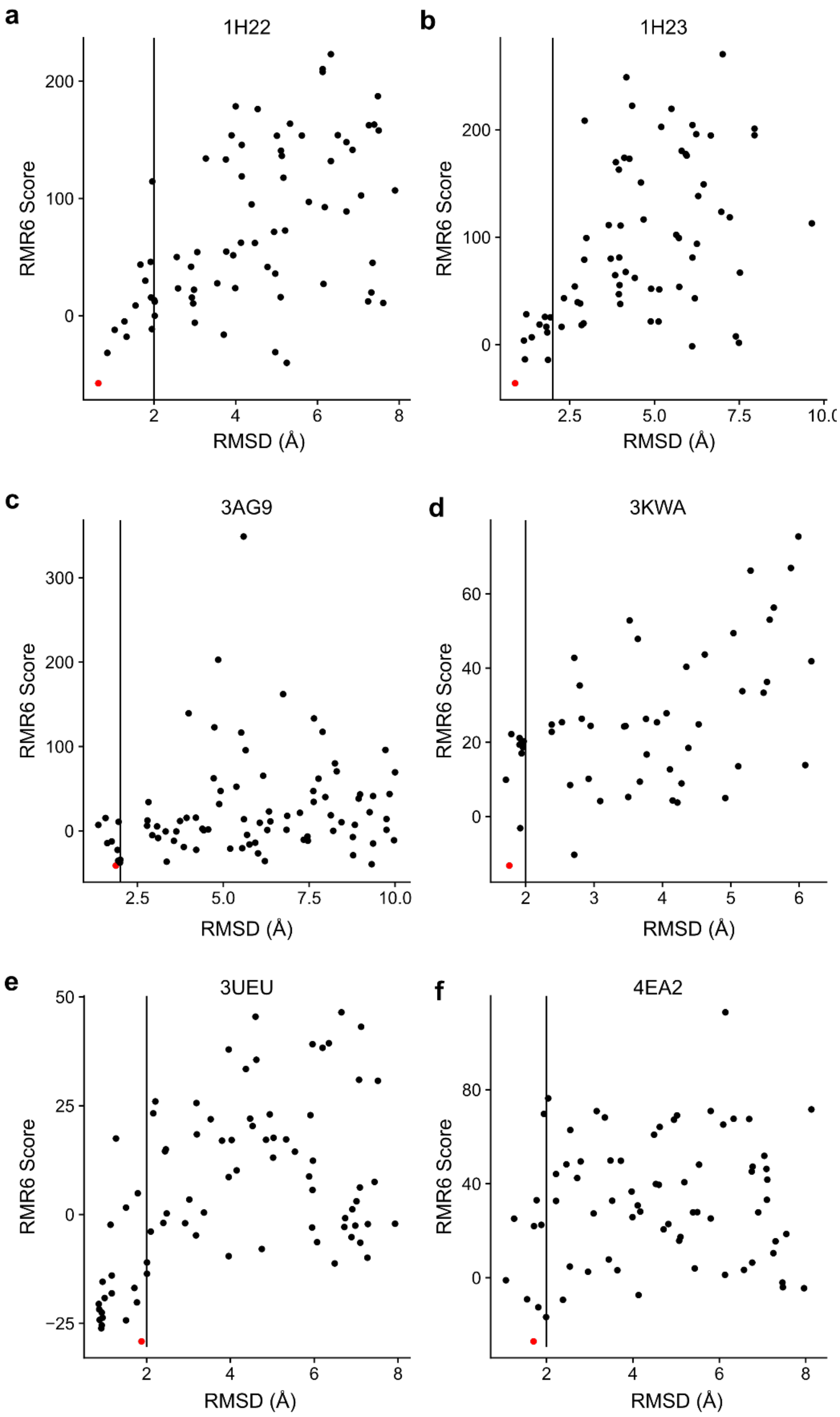


**Figure S5:** Sheep plots for the 6 failure cases detailed in the results and discussion section. In each plot, the RMSD of a CASF-2016 decoy pose is plotted against its RMR6 score where the pose with the lowest RMR6 score is shown in red.

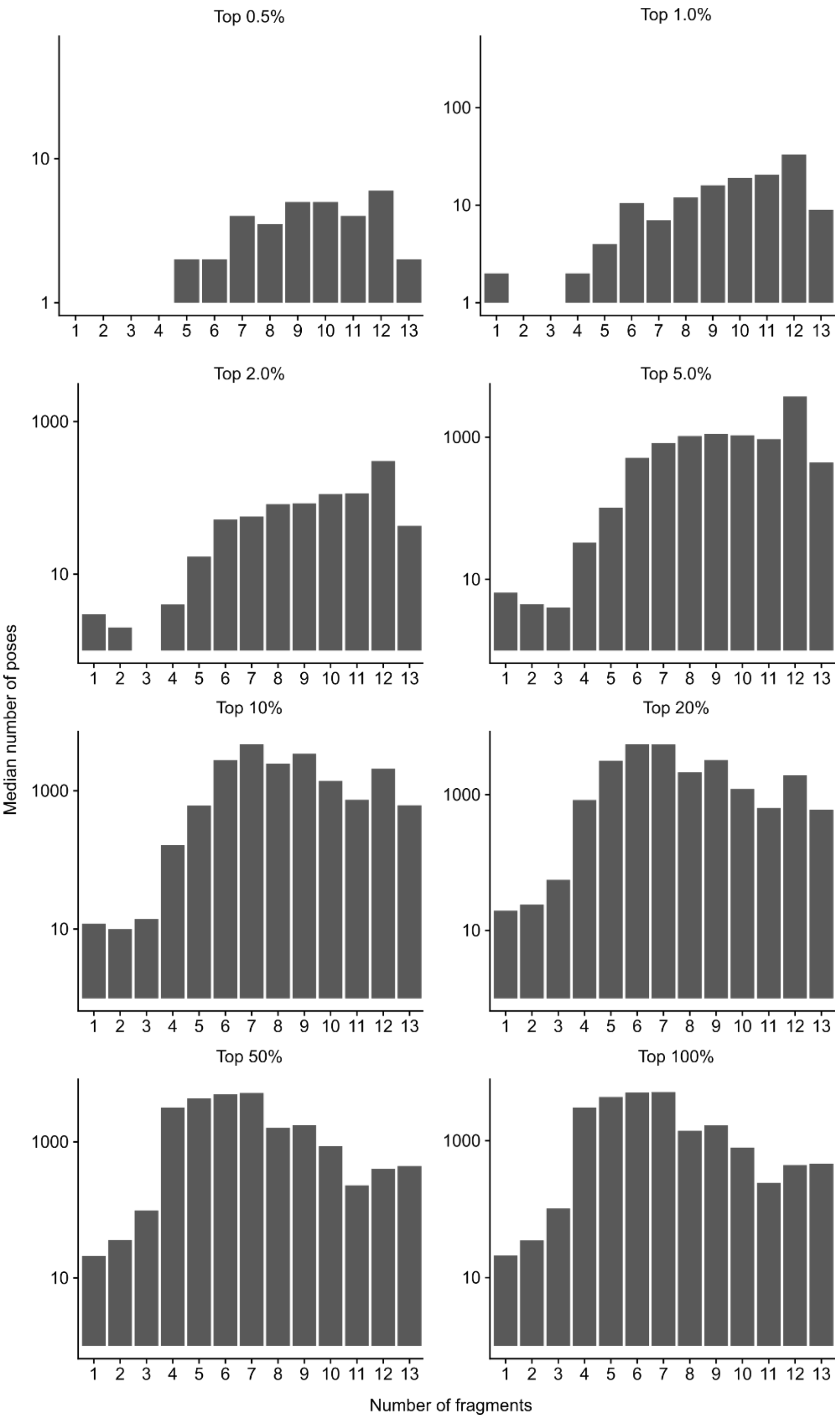


**Figure S6:** Median number of poses generated for ligands containing 1-13 fragments divided by the top percent parameter using semi-flexible algorithm in CANDOCK.

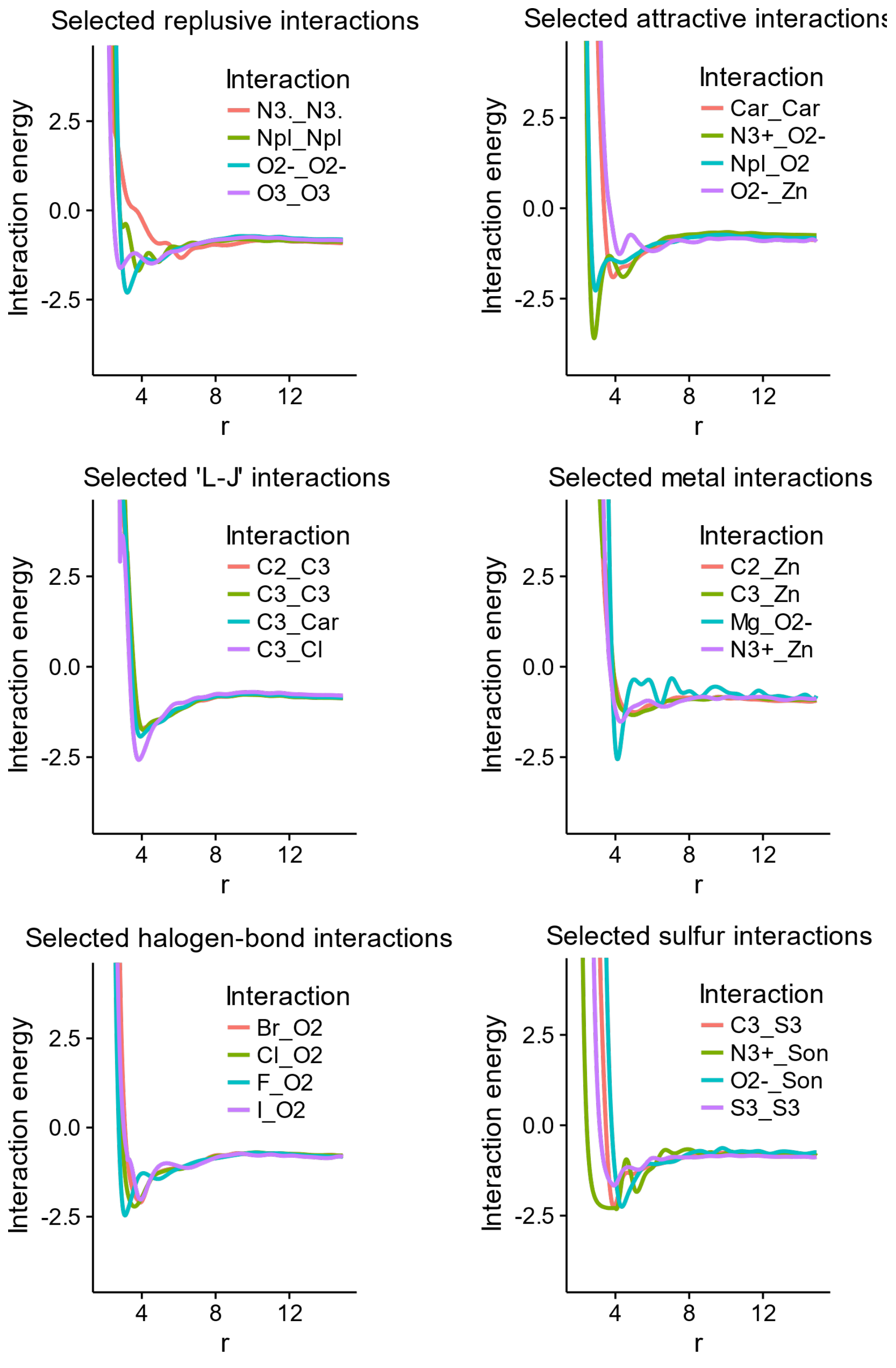


**Figure S7:** Interaction potentials for selected IDATM atom type pairs in the RMC15 objective function. These interactions are selected due to their conventionally repulsive/attractive/neutral nature, or due to their interest in drug discovery.

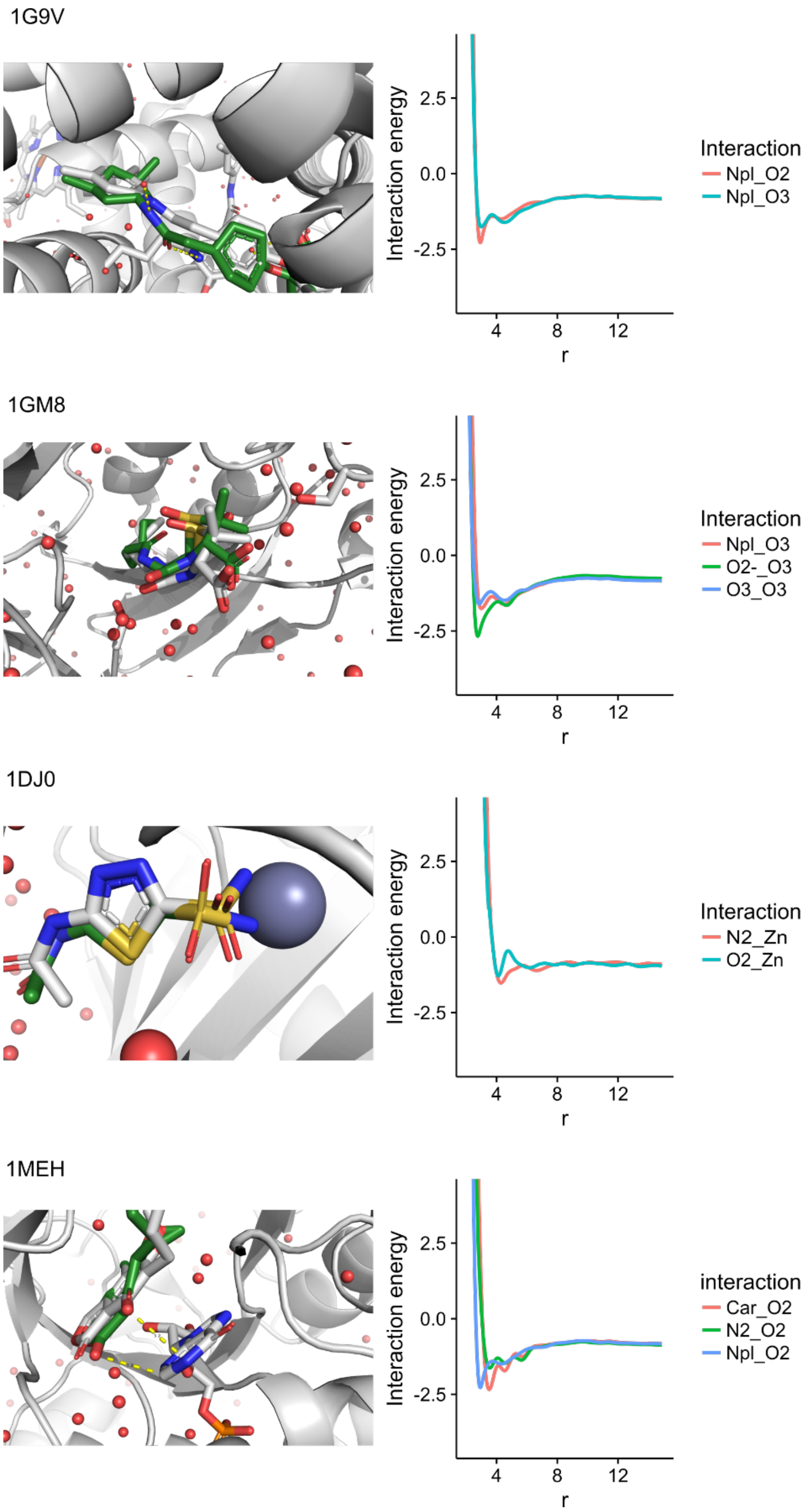


**Figure S8:** Examples where CANDOCK is able to produce a good docking pose where other methods are not able. The best CANDOCK pose is given on the lefthand side of the figure and important interactions between the ligand protein are given on the right.

**Table S1:** Correlations between score and small molecule RMSD calculated and summarized over the entire CASF-2016 benchmarking set.

| | **Rigid protein** | | **Semi-flexible protein** | | **Fully-flexible protein** | |
|---|---|---|---|---|---|---|
| | Average | Median | Average | Median | Average | Median |
| **rmr4** | 0.095 | 0.049 | 0.140035 | 0.079644 | 0.140 | 0.080 |
| **rmr5** | 0.145 | 0.102 | 0.179655 | 0.112795 | 0.180 | 0.113 |
| **rmr6** | 0.176 | 0.115 | 0.18871 | 0.130094 | 0.189 | 0.130 |
| **rmr7** | 0.178 | 0.112 | 0.207084 | 0.139238 | 0.207 | 0.139 |
| **rmr8** | 0.190 | 0.112 | 0.211746 | 0.165313 | 0.212 | 0.165 |
| **rmr9** | 0.189 | 0.123 | 0.214307 | 0.18227 | 0.214 | 0.182 |
| **rmr10** | 0.203 | 0.141 | 0.236163 | 0.212437 | 0.236 | 0.212 |
| **rmr11** | 0.216 | 0.149 | 0.252629 | 0.230083 | 0.253 | 0.230 |
| **rmr12** | 0.225 | 0.165 | 0.262296 | 0.246224 | 0.262 | 0.246 |
| **rmr13** | 0.222 | 0.181 | 0.261284 | 0.256056 | 0.261 | 0.256 |
| **rmr14** | 0.210 | 0.169 | 0.248759 | 0.235548 | 0.249 | 0.236 |
| **rmr15** | 0.183 | 0.121 | 0.217107 | 0.195525 | 0.217 | 0.196 |
| **rmc4** | 0.307 | 0.279 | 0.361189 | 0.359725 | 0.361 | 0.360 |
| **rmc5** | 0.413 | 0.416 | 0.424266 | 0.425995 | 0.424 | 0.426 |
| **rmc6** | 0.459 | 0.473 | 0.462102 | 0.470914 | 0.462 | 0.471 |
| **rmc7** | 0.476 | 0.501 | 0.494936 | 0.497759 | 0.495 | 0.498 |
| **rmc8** | 0.498 | 0.519 | 0.517914 | 0.528137 | 0.518 | 0.528 |
| **rmc9** | 0.523 | 0.542 | 0.537665 | 0.555256 | 0.538 | 0.555 |
| **rmc10** | 0.537 | 0.560 | 0.550387 | 0.558608 | 0.550 | 0.559 |
| **rmc11** | 0.539 | 0.562 | 0.554441 | 0.564875 | 0.554 | 0.565 |
| **rmc12** | 0.546 | 0.571 | 0.561478 | 0.579258 | 0.561 | 0.579 |
| **rmc13** | 0.546 | 0.576 | 0.562225 | 0.584367 | 0.562 | 0.584 |
| **rmc14** | 0.556 | 0.587 | 0.56007 | 0.586941 | 0.560 | 0.587 |
| **rmc15** | 0.559 | 0.581 | 0.558598 | 0.58669 | 0.559 | 0.587 |
| **fmr4** | 0.081 | 0.027 | 0.120917 | 0.070754 | 0.121 | 0.071 |
| **fmr5** | 0.107 | 0.063 | 0.14436 | 0.094075 | 0.144 | 0.094 |
| **fmr6** | 0.125 | 0.064 | 0.154699 | 0.106345 | 0.155 | 0.106 |
| **fmr7** | 0.132 | 0.069 | 0.165974 | 0.108041 | 0.166 | 0.108 |
| **fmr8** | 0.152 | 0.079 | 0.176905 | 0.101773 | 0.177 | 0.102 |
| **fmr9** | 0.149 | 0.078 | 0.183293 | 0.112085 | 0.183 | 0.112 |
| **fmr10** | 0.149 | 0.070 | 0.182985 | 0.11354 | 0.183 | 0.114 |
| **fmr11** | 0.146 | 0.075 | 0.178826 | 0.118613 | 0.179 | 0.119 |
| **fmr12** | 0.135 | 0.062 | 0.164336 | 0.114359 | 0.164 | 0.114 |
| **fmr13** | 0.116 | 0.048 | 0.140079 | 0.083963 | 0.140 | 0.084 |
| **fmr14** | 0.093 | 0.023 | 0.110206 | 0.053444 | 0.110 | 0.053 |
| **fmr15** | 0.062 | -0.014 | 0.070699 | 0.008513 | 0.071 | 0.009 |
| **fmc4** | 0.307 | 0.270 | 0.359536 | 0.353402 | 0.360 | 0.353 |

| | | | | | | |
|---|---|---|---|---|---|---|
| **fmc5** | 0.424 | 0.427 | 0.428663 | 0.429932 | 0.429 | 0.430 |
| **fmc6** | 0.468 | 0.480 | 0.468468 | 0.480115 | 0.468 | 0.480 |
| **fmc7** | 0.486 | 0.510 | 0.501306 | 0.508722 | 0.501 | 0.509 |
| **fmc8** | 0.505 | 0.527 | 0.523356 | 0.530233 | 0.523 | 0.530 |
| **fmc9** | 0.535 | 0.549 | 0.541273 | 0.563213 | 0.541 | 0.563 |
| **fmc10** | 0.539 | 0.560 | 0.551955 | 0.563304 | 0.552 | 0.563 |
| **fmc11** | 0.539 | 0.562 | 0.553678 | 0.569521 | 0.554 | 0.570 |
| **fmc12** | 0.545 | 0.569 | 0.560537 | 0.581049 | 0.561 | 0.581 |
| **fmc13** | 0.546 | 0.580 | 0.561942 | 0.582873 | 0.562 | 0.583 |
| **fmc14** | 0.556 | 0.587 | 0.558451 | 0.586431 | 0.558 | 0.586 |
| **fmc15** | 0.559 | 0.582 | 0.556559 | 0.583299 | 0.557 | 0.583 |
| **rcr4** | 0.050 | -0.015 | 0.115444 | 0.052879 | 0.115 | 0.053 |
| **rcr5** | 0.053 | -0.015 | 0.103434 | 0.042769 | 0.103 | 0.043 |
| **rcr6** | 0.056 | -0.016 | 0.095697 | 0.036087 | 0.096 | 0.036 |
| **rcr7** | 0.047 | -0.018 | 0.08391 | 0.02515 | 0.084 | 0.025 |
| **rcr8** | 0.057 | -0.010 | 0.090313 | 0.022349 | 0.090 | 0.022 |
| **rcr9** | 0.053 | -0.013 | 0.084556 | 0.032991 | 0.085 | 0.033 |
| **rcr10** | 0.039 | -0.025 | 0.06555 | 0.00743 | 0.066 | 0.007 |
| **rcr11** | 0.031 | -0.035 | 0.055276 | 0.009913 | 0.055 | 0.010 |
| **rcr12** | 0.031 | -0.034 | 0.054034 | 0.007386 | 0.054 | 0.007 |
| **rcr13** | 0.038 | -0.031 | 0.064184 | 0.023671 | 0.064 | 0.024 |
| **rcr14** | 0.057 | -0.017 | 0.088855 | 0.036508 | 0.089 | 0.037 |
| **rcr15** | 0.081 | -0.006 | 0.119669 | 0.068518 | 0.120 | 0.069 |
| **rcc4** | 0.067 | 0.001 | 0.118727 | 0.047455 | 0.119 | 0.047 |
| **rcc5** | 0.075 | 0.007 | 0.112314 | 0.046986 | 0.112 | 0.047 |
| **rcc6** | 0.073 | 0.013 | 0.099047 | 0.039596 | 0.099 | 0.040 |
| **rcc7** | 0.056 | -0.005 | 0.078413 | 0.021166 | 0.078 | 0.021 |
| **rcc8** | 0.051 | -0.012 | 0.066573 | -1.97E-04 | 0.067 | 0.000 |
| **rcc9** | 0.036 | -0.028 | 0.047623 | -0.0104 | 0.048 | -0.010 |
| **rcc10** | 0.020 | -0.053 | 0.024173 | -0.02832 | 0.024 | -0.028 |
| **rcc11** | 0.011 | -0.073 | 0.013487 | -0.04264 | 0.013 | -0.043 |
| **rcc12** | 0.013 | -0.072 | 0.013389 | -0.04284 | 0.013 | -0.043 |
| **rcc13** | 0.021 | -0.061 | 0.02453 | -0.03049 | 0.025 | -0.030 |
| **rcc14** | 0.040 | -0.044 | 0.049441 | -0.00867 | 0.049 | -0.009 |
| **rcc15** | 0.065 | -0.010 | 0.080644 | 0.019915 | 0.081 | 0.020 |
| **fcr4** | 0.050 | -0.015 | 0.114892 | 0.061528 | 0.115 | 0.062 |
| **fcr5** | 0.062 | -0.002 | 0.1126 | 0.060021 | 0.113 | 0.060 |
| **fcr6** | 0.071 | -0.001 | 0.116733 | 0.055982 | 0.117 | 0.056 |
| **fcr7** | 0.074 | 0.003 | 0.122048 | 0.056749 | 0.122 | 0.057 |
| **fcr8** | 0.084 | 0.014 | 0.129655 | 0.070466 | 0.130 | 0.070 |
| **fcr9** | 0.086 | 0.007 | 0.131851 | 0.072608 | 0.132 | 0.073 |

| | | | | | | |
|---|---|---|---|---|---|---|
| **fcr10** | 0.091 | 0.011 | 0.137708 | 0.071559 | 0.138 | 0.072 |
| **fcr11** | 0.102 | 0.033 | 0.153892 | 0.105595 | 0.154 | 0.106 |
| **fcr12** | 0.116 | 0.047 | 0.171489 | 0.114962 | 0.171 | 0.115 |
| **fcr13** | 0.133 | 0.063 | 0.193219 | 0.14408 | 0.193 | 0.144 |
| **fcr14** | 0.157 | 0.091 | 0.221047 | 0.180568 | 0.221 | 0.181 |
| **fcr15** | 0.177 | 0.116 | 0.243947 | 0.208698 | 0.244 | 0.209 |
| **fcc4** | 0.063 | -0.006 | 0.112292 | 0.05447 | 0.112 | 0.054 |
| **fcc5** | 0.075 | 0.002 | 0.110992 | 0.055188 | 0.111 | 0.055 |
| **fcc6** | 0.084 | 0.019 | 0.116043 | 0.056761 | 0.116 | 0.057 |
| **fcc7** | 0.088 | 0.026 | 0.12244 | 0.058866 | 0.122 | 0.059 |
| **fcc8** | 0.098 | 0.021 | 0.130706 | 0.063725 | 0.131 | 0.064 |
| **fcc9** | 0.101 | 0.029 | 0.134228 | 0.069798 | 0.134 | 0.070 |
| **fcc10** | 0.105 | 0.036 | 0.140838 | 0.075632 | 0.141 | 0.076 |
| **fcc11** | 0.117 | 0.053 | 0.157337 | 0.100401 | 0.157 | 0.100 |
| **fcc12** | 0.132 | 0.071 | 0.175309 | 0.121073 | 0.175 | 0.121 |
| **fcc13** | 0.146 | 0.086 | 0.192998 | 0.13832 | 0.193 | 0.138 |
| **fcc14** | 0.163 | 0.107 | 0.214633 | 0.165242 | 0.215 | 0.165 |
| **fcc15** | 0.179 | 0.126 | 0.232661 | 0.192197 | 0.233 | 0.192 |

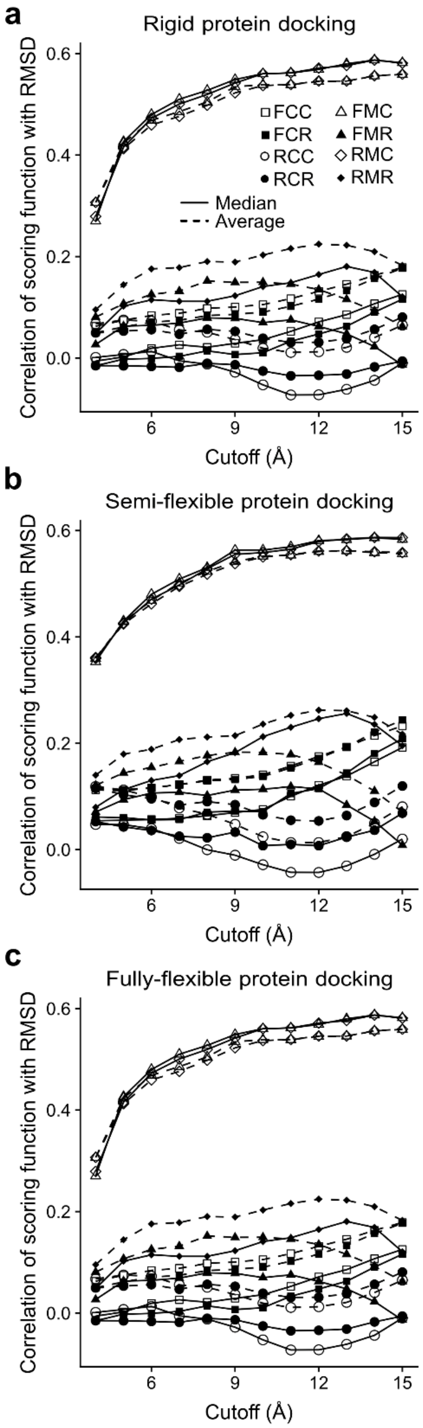


**Figure S9:** Correlations between score and the RMSD of pose from the crystal pose for rigid protein (**a**), semi-flexible protein (**b**), and fully flexible proteins (**c**). These plots show that the RMC and FMC scoring function families produce scores that best correlate with the RMSD of a crystal pose and that as the cutoff of the scoring function increase, so does the correlation with RMSD for these scoring functions.

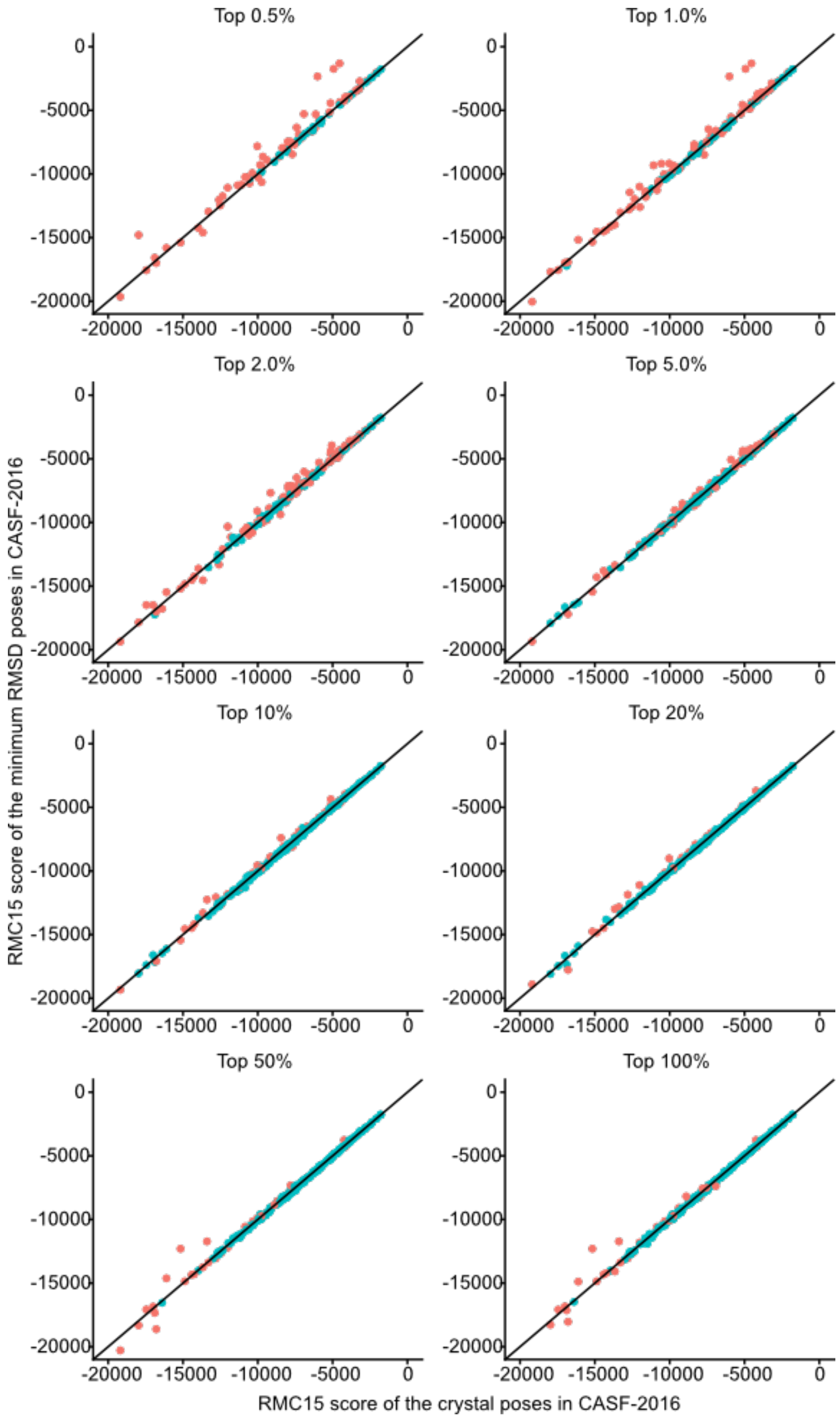


**Figure S10:** Correlations between the RMC15 scores of the crystal pose and the pose with the RMC15 score of the lowest RMSD are shown for all eight top percent values. Poses within 2.0 Å of the crystal pose are shown in blue (successful runs) while poses greater than 2.0 Å are shown in red.

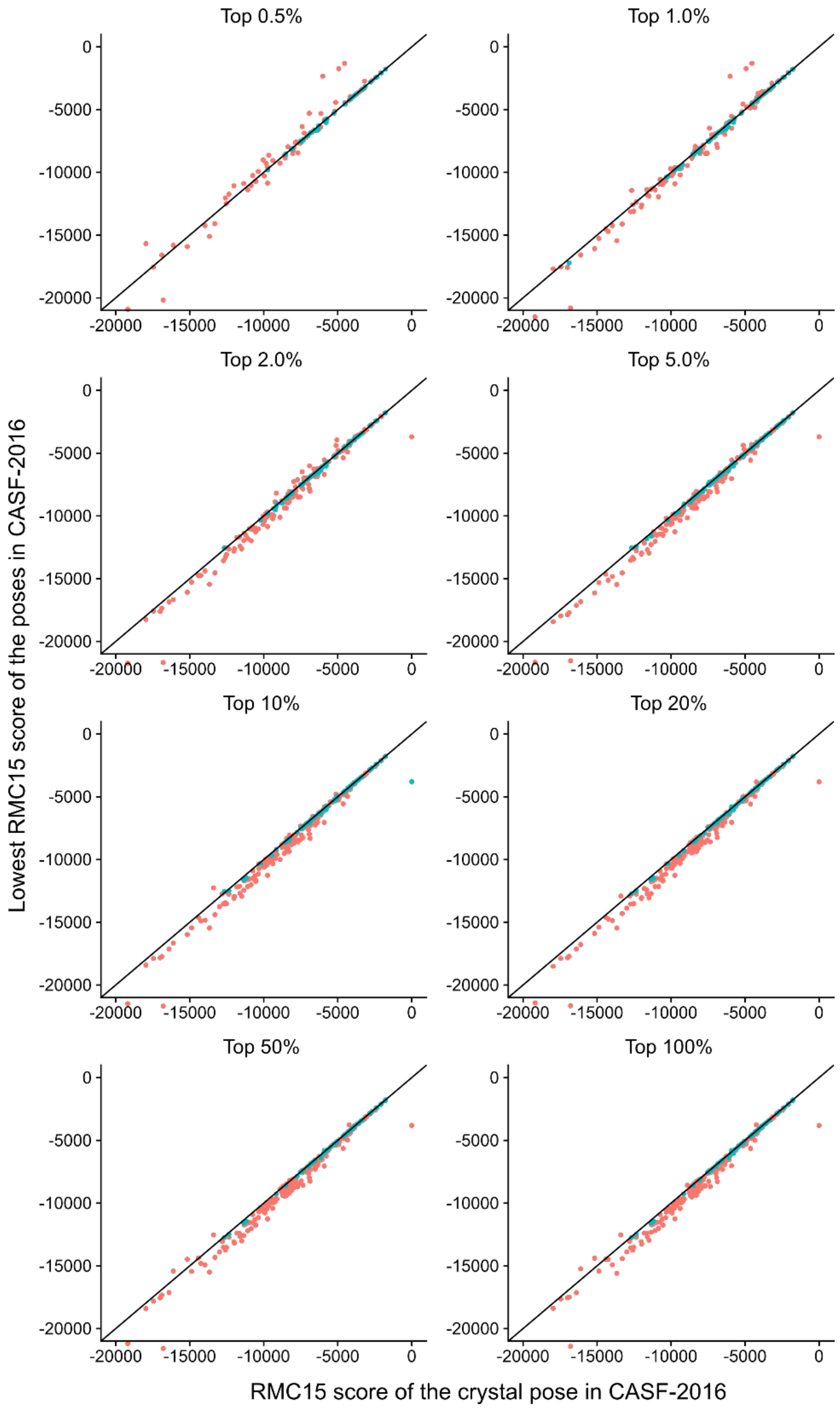


**Figure S11:** The RMC15 score for the pose selected by RMR6 obtained for each cocrystal is plotted against the RMC15 score of the crystal pose. Poses within 2.0 Å of the crystal pose are shown in blue (successful runs) while poses greater than 2.0 Å are shown in red. Here it is shown that the successful poses occur only on the y=x line, while the unsuccessful poses cluster above this line. This indicates that further minimization with RMC15 may improve the RMR6 selection rate.

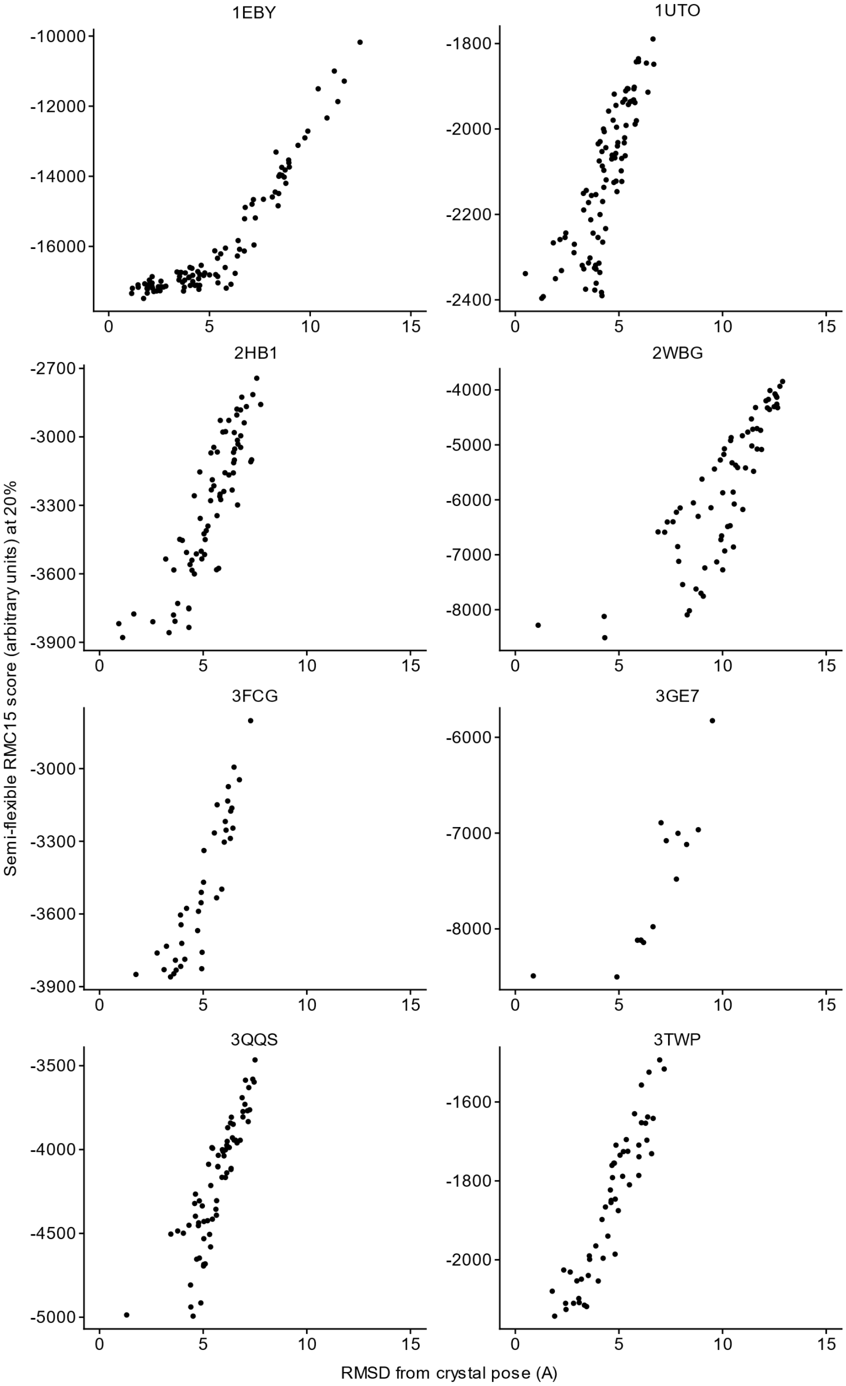


**Figure S12:** Plots of the RMC15 score of all poses produced by CANDOCK for selected proteins in CASF-2016 versus the RMSD of the pose. In all plots, the RMSD ranges from 1Å to 15Å. The poses were obtained using the semi flexible method at a 'Top Percent' value of 20%. The trend seen for these proteins is as one decreases the RMC15 score, the RMSD of the predicted ligand also decreases. Therefore, the use of an objective function derived from the RMC15 scoring function to minimize the ligand in the binding pocket is justified.

**Table S2:** Pearson correlations for all ligands in CASF-2016 using various scoring functions to select the representative pose for the docked (semi-flexible) protein-ligand complex and rank the activity of the ligand versus other ligands for the same protein. The orange highlighted cell values are plotted in **Figure S14.**

| Selector: | RMSD | | RMR6 | | RMC15 | |
|---|---|---|---|---|---|---|
| **Ranker** | **RMR6** | **RMC15** | **RMR6** | **RMC15** | **RMR6** | **RMC15** |
| **3-DEHYDROQUINATE DEHYDRATASE** | -0.716 | -0.876 | -0.852 | -0.875 | 0.591 | -0.874 |
| **ACETYLCHOLINE RECEPTOR** | 0.335 | 0.339 | 0.476 | 0.350 | 0.772 | 0.326 |
| **ACETYLCHOLINE-BINDING PROTEIN** | 0.482 | -0.193 | -0.004 | -0.195 | 0.554 | -0.212 |
| **ACHE** | -0.269 | -0.664 | -0.474 | -0.688 | -0.300 | -0.652 |
| **ALPHA-L-FUCOSIDASE** | -0.692 | -0.372 | -0.606 | -0.307 | -0.666 | -0.304 |
| **ALPHA-MANNOSIDASE II** | -0.271 | -0.581 | 0.575 | -0.580 | 0.855 | -0.599 |
| **ANDROGEN RECEPTOR** | -0.919 | 0.734 | -0.738 | 0.730 | -0.776 | 0.736 |
| **TrpD** | 0.652 | -0.905 | -0.328 | -0.832 | 0.539 | -0.920 |
| **BETA-GLUCOSIDASE A** | 0.751 | 0.140 | -0.337 | 0.119 | -0.953 | 0.147 |
| **BETA-LACTAMASE** | -0.495 | -0.894 | -0.681 | -0.908 | 0.835 | -0.886 |
| **BETA-LACTOGLOBULIN** | -0.981 | -0.991 | -0.978 | -0.997 | -0.959 | -0.993 |
| **BETA-SECRETASE 1** | 0.695 | -0.116 | -0.210 | -0.370 | 0.673 | -0.121 |
| **BROMODOMAIN-CONTAINING PROTEIN 4** | -0.644 | -0.981 | -0.579 | -0.988 | 0.393 | -0.955 |
| **CAMP-DEPENDENT PROTEIN KINASE** | 0.696 | -0.905 | -0.619 | -0.996 | 0.785 | -0.864 |
| **CARBONIC ANHYDRASE 2** | -0.694 | -0.883 | 0.770 | -0.770 | 0.976 | -0.856 |
| **CATECHOL O-METHYLTRANSFERASE** | -0.858 | -0.870 | -0.749 | -0.839 | -0.687 | -0.781 |
| **CELL DIVISION PROTEIN KINASE 2** | -0.800 | -0.899 | 0.713 | -0.918 | 0.969 | -0.879 |

| | | | | | | |
|---|---|---|---|---|---|---|
| **CELLULAR TUMOR ANTIGEN P53** | -0.739 | -0.719 | 0.796 | -0.719 | 0.925 | -0.648 |
| **CGMP 3',5'-CYCLIC PHOSPHODIESTERASE** | -0.649 | 0.052 | -0.138 | 0.103 | -0.711 | 0.074 |
| **CHITINASE A** | -0.915 | -0.725 | -0.833 | -0.725 | -0.704 | -0.682 |
| **COAGULATION FACTOR XA** | -0.963 | -0.753 | -0.992 | -0.671 | 0.560 | -0.596 |
| **FACTOR XI** | -0.805 | -0.916 | -0.864 | -0.885 | 0.989 | -0.887 |
| **DEHYDROSQUALENE SYNTHASE** | -0.353 | -0.439 | -0.727 | -0.468 | 0.173 | -0.432 |
| **ENDOTHIAPEPSIN** | -0.975 | -0.993 | -0.903 | -0.992 | 0.971 | -0.989 |
| **ESTROGEN RECEPTOR** | -0.843 | 0.764 | 0.501 | 0.764 | -0.293 | 0.761 |
| **GLUTAMATE RECEPTOR 2** | -0.814 | -0.457 | -0.741 | -0.458 | -0.854 | -0.454 |
| **GLUTAMATE RECEPTOR, IONOTROPIC KAINATE 1** | -0.977 | -0.646 | -0.702 | -0.632 | 0.872 | -0.577 |
| **GLYCOGEN PHOSPHORYLASE** | 0.716 | 0.383 | -0.186 | 0.407 | -0.833 | 0.341 |
| **HEAT SHOCK PROTEIN HSP82** | -0.418 | 0.112 | -0.574 | 0.105 | -0.771 | 0.166 |
| **HEAT SHOCK PROTEIN HSP90-ALPHA** | -0.728 | -0.879 | -0.832 | -0.882 | 0.778 | -0.873 |
| **HIV-1 INTEGRASE** | -0.843 | -0.954 | -0.954 | -0.960 | 0.935 | -0.952 |
| **HIV-1 PROTEASE** | -0.916 | -0.573 | -0.907 | -0.468 | -0.926 | -0.541 |
| **(MMP-1)** | -0.680 | -0.815 | -0.801 | -0.808 | 0.966 | -0.796 |
| **MITOGEN-ACTIVATED PROTEIN KINASE 14** | -0.680 | -0.902 | -0.691 | -0.903 | 0.916 | -0.900 |
| **MTA/SAH NUCLEOSIDASE** | 0.601 | 0.203 | 0.601 | 0.203 | 0.657 | 0.206 |
| **O-GLCNACASE BT_4395** | -0.974 | -0.128 | -0.471 | -0.129 | -0.392 | -0.074 |
| **PANTOTHENATE SYNTHETASE** | -0.873 | -0.960 | -0.764 | -0.961 | 0.835 | -0.962 |
| **PPARG** | -0.974 | -0.992 | -0.971 | -0.989 | 0.902 | -0.983 |

| | | | | | | |
|---|---|---|---|---|---|---|
| **PROTEIN-TYROSINE PHOSPHATASE 1B** | 0.304 | -0.947 | -0.080 | -0.805 | 0.759 | -0.874 |
| **QUEUINE TRNA-RIBOSYLTRANSFERASE** | -0.875 | -0.697 | -0.875 | -0.697 | -0.676 | -0.703 |
| **RIBONUCLEASE A** | -0.701 | -0.842 | -0.908 | -0.849 | 0.708 | -0.836 |
| **RNA-DIRECTED RNA POLYMERASE** | -0.794 | -0.831 | -0.740 | -0.828 | 0.981 | -0.856 |
| **SERINE/THREONINE-PROTEIN KINASE 6** | -0.093 | -0.706 | 0.091 | -0.885 | -0.603 | -0.619 |
| **CHK1** | 0.509 | 0.545 | -0.204 | 0.637 | 0.963 | 0.386 |
| **PIM-1** | 0.656 | 0.447 | 0.423 | 0.436 | -0.568 | 0.424 |
| **TANKYRASE-2** | -0.918 | -0.854 | -0.970 | -0.881 | 0.491 | -0.825 |
| **THERMOLYSIN** | -0.610 | 0.178 | -0.546 | -0.272 | -0.700 | 0.149 |
| **THROMBIN** | -0.798 | 0.154 | 0.473 | 0.188 | 0.562 | 0.155 |
| **TRANSCRIPTION ELONGATION FACTOR B POLYPEPTIDE 2** | 0.789 | -0.772 | 0.784 | -0.756 | 0.908 | -0.770 |
| **TRANSPORTER** | -0.376 | 0.497 | -0.284 | 0.509 | -0.352 | 0.510 |
| **TRYPSIN BETA** | -0.905 | -0.805 | -0.927 | -0.804 | 0.653 | -0.769 |
| **TYROSINE-PROTEIN KINASE ABL1** | 0.929 | -0.119 | 0.187 | 0.093 | 0.847 | -0.221 |
| **TYROSINE-PROTEIN KINASE ITK/TSK** | 0.879 | 0.749 | 0.183 | 0.747 | -0.786 | 0.755 |
| **TYROSINE-PROTEIN KINASE JAK1** | -0.552 | -0.093 | -0.292 | -0.154 | 0.234 | -0.079 |
| **TYROSINE-PROTEIN KINASE JAK2** | -0.653 | -0.883 | -0.718 | -0.876 | 0.759 | -0.931 |
| **UROKINASE-TYPE PLASMINOGEN ACTIVATOR** | -0.909 | -0.927 | -0.908 | -0.922 | 0.882 | -0.924 |

**Table S3:** Spearman correlations for all ligands in CASF-2016 using various scoring functions to select the representative pose for the docked (semi-flexible) protein-ligand complex and rank the activity of the ligand versus other ligands for the same protein. The orange highlighted cell values are plotted in **Figure S14.**

| Selector | RMSD | | RMR6 | | RMC15 | |
|---|---|---|---|---|---|---|
| **Ranker** | **RMR6** | **RMC15** | **RMR6** | **RMC15** | **RMR6** | **RMC15** |
| **3-DEHYDROQUINATE DEHYDRATASE** | -0.400 | -0.600 | -1.000 | -0.600 | 0.700 | -0.600 |
| **ACETYLCHOLINE RECEPTOR** | 0.400 | 0.100 | 0.600 | 0.100 | 0.600 | 0.100 |
| **ACETYLCHOLINE-BINDING PROTEIN** | 0.500 | -0.300 | -0.600 | -0.300 | 0.700 | -0.300 |
| **ACHE** | -0.500 | -0.700 | -0.400 | -0.700 | -0.300 | -0.700 |
| **ALPHA-L-FUCOSIDASE** | -0.700 | 0.400 | -0.700 | 0.400 | -0.700 | 0.400 |
| **ALPHA-MANNOSIDASE II** | -0.300 | -0.300 | 0.300 | -0.300 | 0.700 | -0.300 |
| **ANDROGEN RECEPTOR** | -0.700 | 0.900 | -0.600 | 0.900 | -1.000 | 0.900 |
| **TrpD** | 0.500 | -1.000 | -0.500 | -0.900 | 0.300 | -1.000 |
| **BETA-GLUCOSIDASE A** | 0.700 | 0.300 | -0.400 | 0.300 | -1.000 | 0.300 |
| **BETA-LACTAMASE** | -0.455 | -0.782 | -0.745 | -0.879 | 0.758 | -0.782 |
| **BETA-LACTOGLOBULIN** | -1.000 | -1.000 | -1.000 | -1.000 | -1.000 | -1.000 |
| **BETA-SECRETASE 1** | 0.500 | 0.000 | -0.300 | -0.400 | 0.700 | 0.000 |
| **BROMODOMAIN-CONTAINING PROTEIN 4** | -0.700 | -0.900 | -0.600 | -1.000 | 0.200 | -0.900 |
| **CAMP-DEPENDENT PROTEIN KINASE** | 0.700 | -1.000 | -0.400 | -1.000 | 0.600 | -1.000 |
| **CARBONIC ANHYDRASE 2** | -0.600 | -0.800 | 0.700 | -0.800 | 0.900 | -0.800 |
| **CATECHOL O-METHYLTRANSFERASE** | -0.800 | -0.900 | -0.700 | -0.900 | -0.900 | -0.900 |
| **CELL DIVISION PROTEIN KINASE 2** | -0.600 | -0.700 | 0.500 | -0.700 | 0.900 | -0.700 |

| | | | | | | |
|---|---|---|---|---|---|---|
| **CELLULAR TUMOR ANTIGEN P53** | -0.900 | -0.700 | 0.900 | -0.700 | 0.900 | -0.400 |
| **CGMP 3',5'-CYCLIC PHOSPHODIESTERASE** | -0.700 | 0.000 | -0.300 | 0.000 | -0.700 | 0.000 |
| **CHITINASE A** | -0.900 | -0.700 | -0.900 | -0.700 | -0.800 | -0.700 |
| **FACTOR XA** | -0.900 | -0.900 | -1.000 | -0.900 | 0.700 | -0.900 |
| **FACTOR XI** | -0.700 | -0.600 | -0.700 | -0.300 | 0.900 | -0.300 |
| **DEHYDROSQUALENE SYNTHASE** | -0.600 | -0.100 | -0.800 | -0.200 | 0.500 | -0.600 |
| **ENDOTHIAPEPSIN** | -1.000 | -1.000 | -0.900 | -0.900 | 1.000 | -0.900 |
| **ESTROGEN RECEPTOR** | -0.800 | 0.300 | 0.300 | 0.300 | -0.600 | 0.300 |
| **GLUTAMATE RECEPTOR 2** | -0.700 | -0.700 | -0.700 | -0.700 | -0.900 | -0.700 |
| **GLUTAMATE RECEPTOR, IONOTROPIC KAINATE 1** | -1.000 | -0.700 | -0.700 | -0.700 | 0.900 | -0.700 |
| **GLYCOGEN PHOSPHORYLASE** | 0.900 | 0.300 | 0.200 | 0.300 | -0.800 | 0.300 |
| **HEAT SHOCK PROTEIN HSP82** | -0.500 | 0.400 | -0.600 | 0.400 | -0.800 | 0.400 |
| **HEAT SHOCK PROTEIN HSP90-ALPHA** | -0.600 | -0.800 | -0.700 | -0.800 | 0.900 | -0.800 |
| **HIV-1 INTEGRASE** | -0.900 | -1.000 | -0.900 | -1.000 | 0.700 | -1.000 |
| **HIV-1 PROTEASE** | -0.900 | -0.700 | -0.900 | -0.800 | -1.000 | -0.600 |
| **MMP-12** | -0.900 | -0.900 | -0.900 | -0.900 | 0.900 | -0.900 |
| **MITOGEN-ACTIVATED PROTEIN KINASE 14** | -0.700 | -0.900 | -0.700 | -1.000 | 0.900 | -0.900 |
| **MTA/SAH NUCLEOSIDASE** | 0.700 | -0.100 | 0.700 | -0.100 | 0.600 | -0.100 |
| **O-GLCNACASE BT_4395** | -0.900 | -0.600 | -0.500 | -0.600 | -0.600 | -0.600 |
| **PANTOTHENATE SYNTHETASE** | -0.900 | -1.000 | -0.900 | -1.000 | 0.800 | -1.000 |
| **PPARG** | -0.700 | -1.000 | -1.000 | -1.000 | 1.000 | -1.000 |

| | | | | | | |
|---|---|---|---|---|---|---|
| **PROTEIN-TYROSINE PHOSPHATASE 1B** | 0.400 | -1.000 | -0.300 | -0.700 | 0.700 | -0.900 |
| **QUEUINE TRNA-RIBOSYLTRANSFERASE** | -0.700 | -0.600 | -0.700 | -0.600 | -0.600 | -0.600 |
| **RIBONUCLEASE A** | -0.700 | -0.700 | -0.900 | -0.700 | 0.400 | -0.700 |
| **RNA-DIRECTED RNA POLYMERASE** | -0.600 | -1.000 | -0.900 | -1.000 | 1.000 | -1.000 |
| **SERINE/THREONINE-PROTEIN KINASE 6** | -0.500 | -0.900 | -0.500 | -0.900 | -0.600 | -0.700 |
| **CHK1** | 0.600 | 0.600 | -0.400 | 0.600 | 1.000 | 0.600 |
| **PIM-1** | 0.300 | 0.300 | 0.200 | 0.300 | -0.300 | 0.300 |
| **TANKYRASE-2** | -1.000 | -0.800 | -0.900 | -0.900 | 0.500 | -0.800 |
| **THERMOLYSIN** | -0.600 | 0.000 | -0.600 | -0.400 | -0.400 | 0.000 |
| **THROMBIN** | -0.800 | 0.400 | 0.300 | 0.400 | 0.500 | 0.200 |
| **TCEB2** | 0.700 | -0.700 | 0.800 | -0.700 | 0.800 | -0.700 |
| **TRANSPORTER** | -0.500 | 0.700 | -0.300 | 0.700 | -0.600 | 0.700 |
| **TRYPSIN BETA** | -0.900 | -0.800 | -0.900 | -0.800 | 0.600 | -0.800 |
| **TYROSINE-PROTEIN KINASE ABL1** | 1.000 | -0.300 | -0.400 | -0.300 | 0.900 | -0.300 |
| **TYROSINE-PROTEIN KINASE ITK/TSK** | 0.900 | 0.700 | -0.100 | 0.700 | -0.400 | 0.500 |
| **TYROSINE-PROTEIN KINASE JAK1** | -0.600 | -0.300 | -0.600 | -0.300 | 0.500 | -0.300 |
| **TYROSINE-PROTEIN KINASE JAK2** | -0.700 | -0.900 | -0.900 | -0.900 | 0.700 | -0.900 |
| **UROKINASE-TYPE PLASMINOGEN ACTIVATOR** | -0.900 | -0.900 | -0.900 | -0.900 | 1.000 | -0.900 |

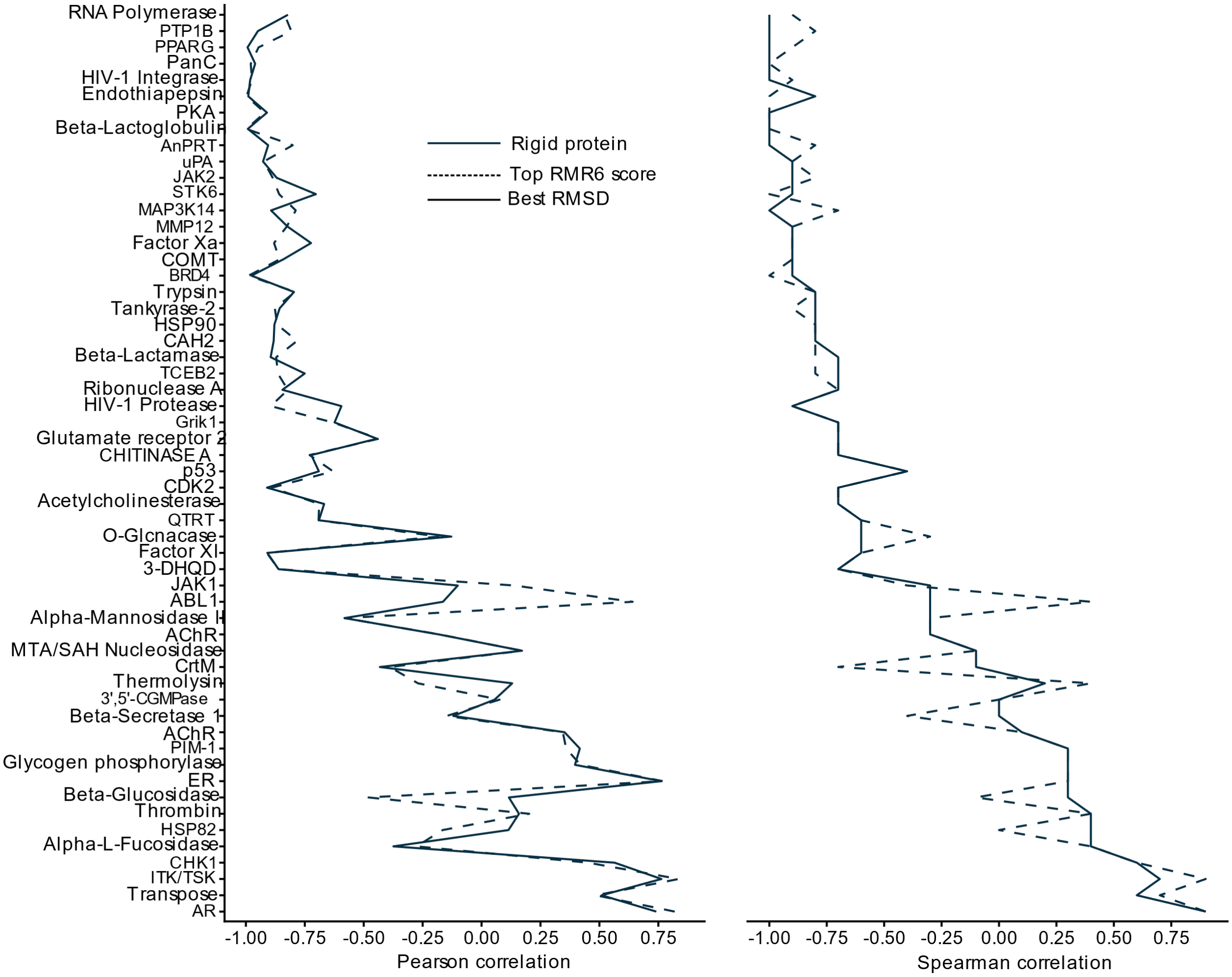


**Figure S13:** Rigid protein docking correlations between the RMC15 score and the measured pKd/pKi of the compounds in CASF-2016 for each protein target. A negative correlation is expected as a decrease in score (an estimation of free energy change) should result in an increase in the negative log of the binding coefficient. The representative docked ligand pose for ranking was selected with either the lowest RMSD or the best RMR6 score criterion.

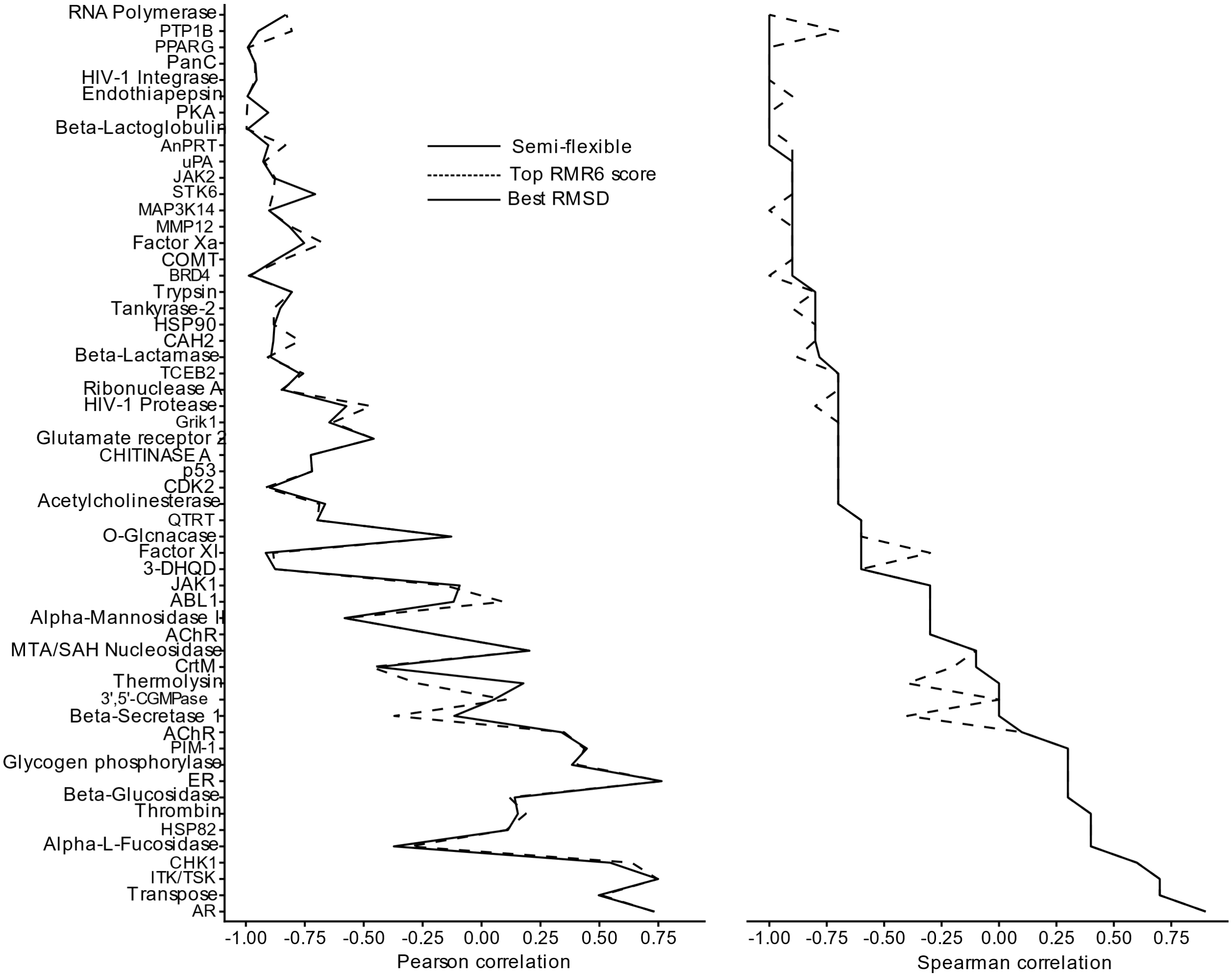


**Figure S14:** Semi-flexible protein docking correlations between the RMC15 score and the measured pKd/pKi of the compounds in CASF-2016 for each protein target. The representative docked ligand pose for ranking was selected with either the lowest RMSD or the best RMR6 score (our best selector) criterion.

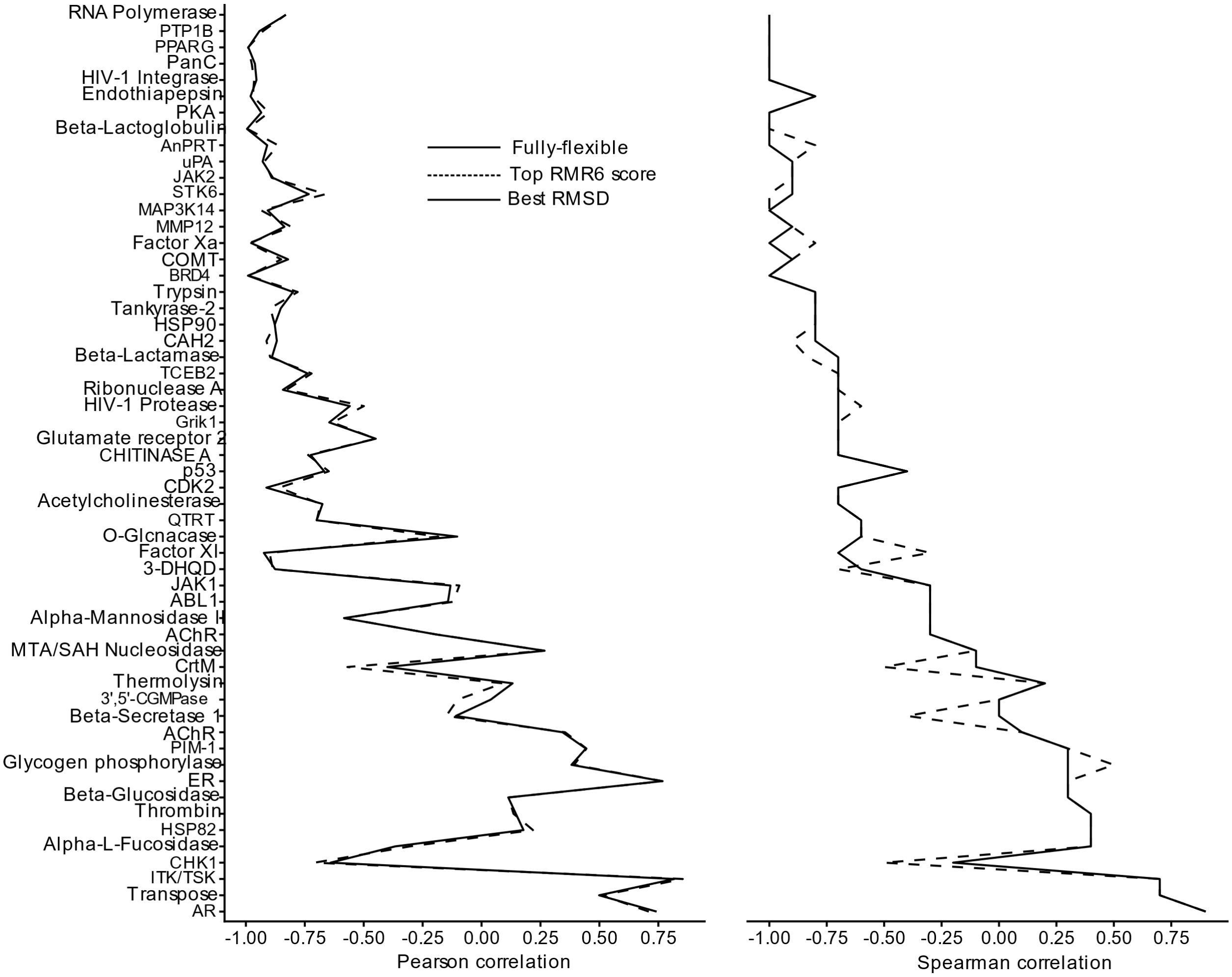


**Figure S15:** Fully flexible protein docking correlations between the RMC15 score and the measured pKd/pKi of the compounds in CASF-2016 for each protein target. The representative docked ligand pose for ranking was selected with either the lowest RMSD or the best RMR6 score (our best selector) criterion.

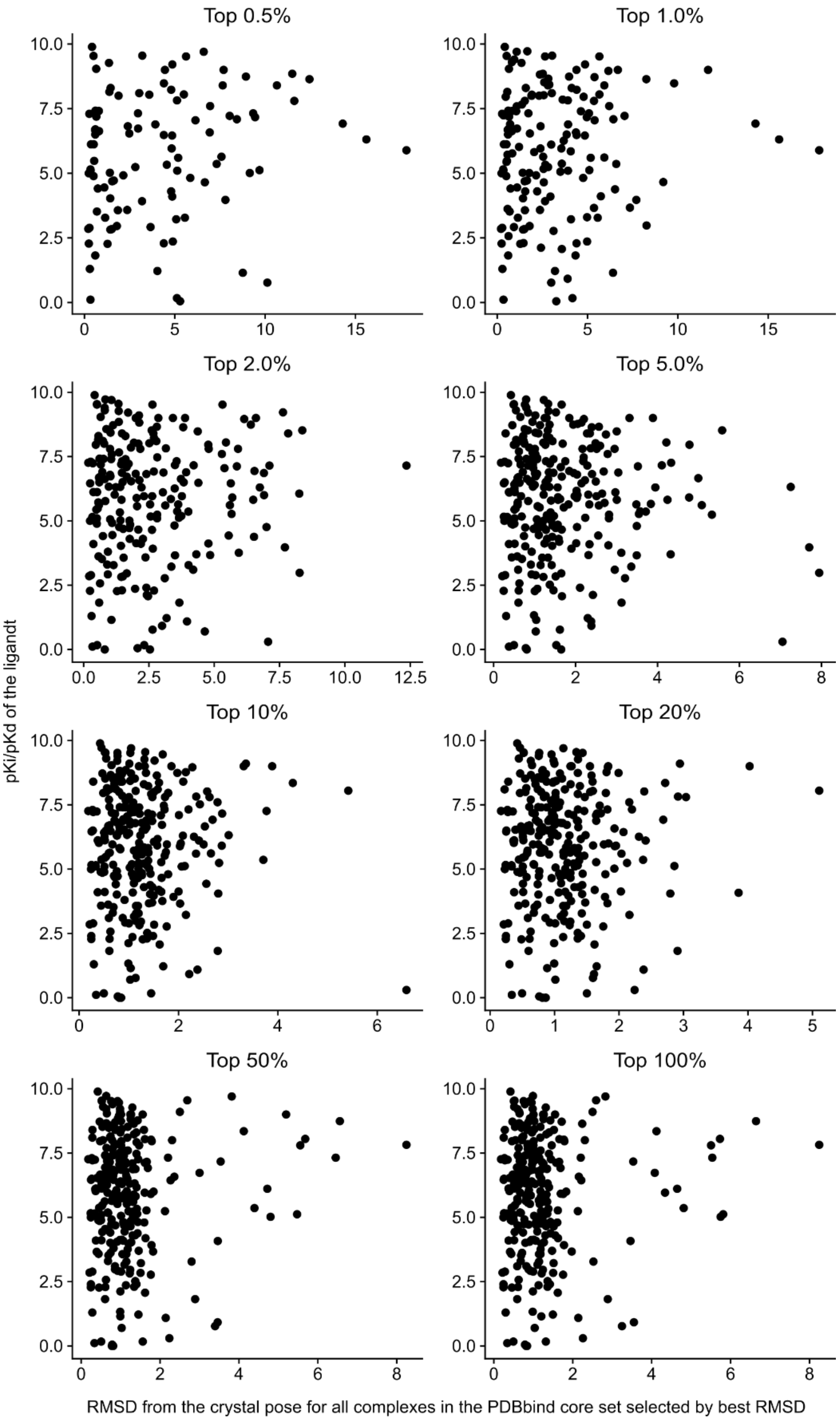


**Figure S16:** Plots of the ligand pKi/pKd against the best pose RMSD obtained using the semi-flexible method broken down by the 'Top Percent' parameter. These plots show there is no relationship between RMSD and pKi.

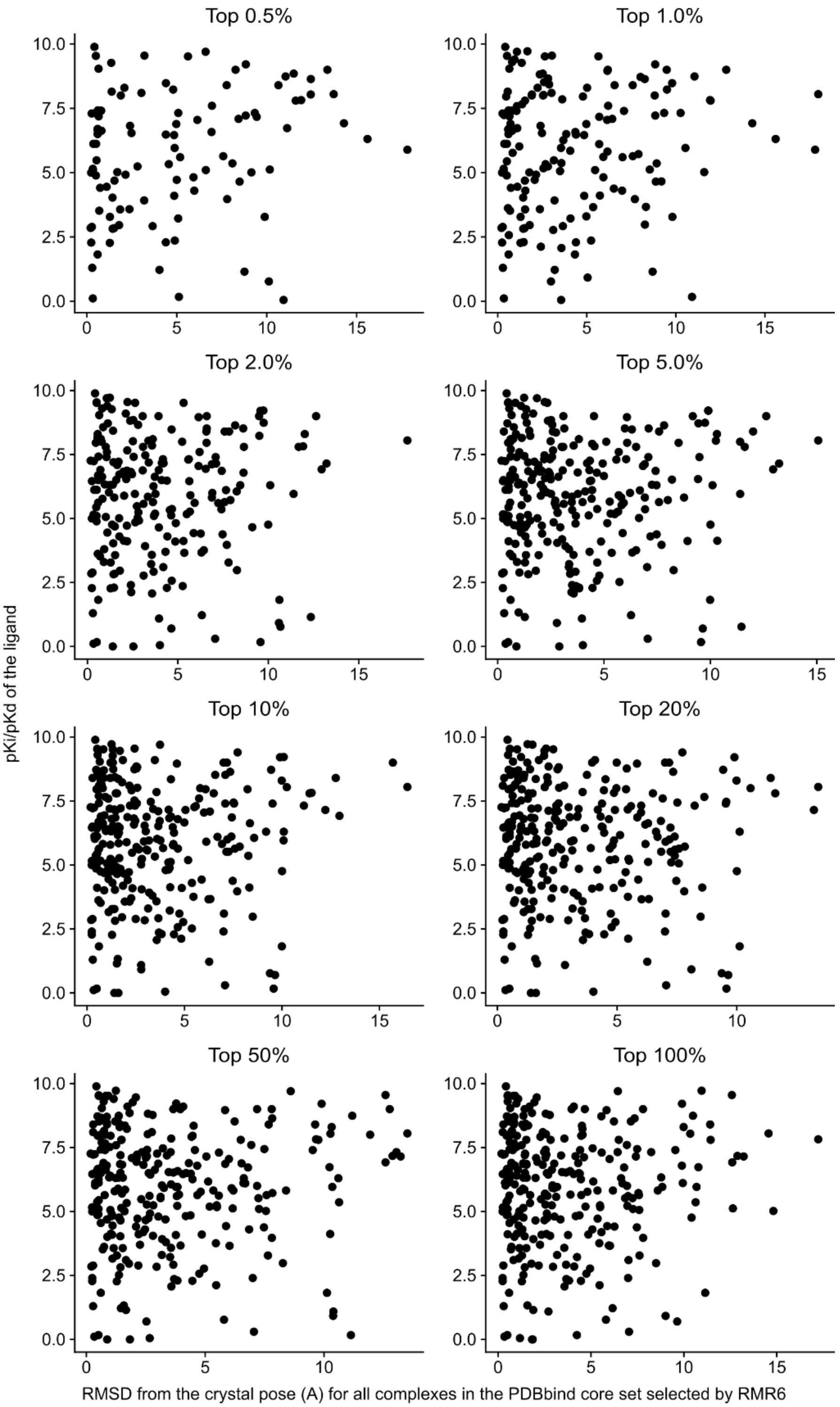


**Figure S17:** Plots of the ligand pKi/pKd against the RMR selected pose RMSD obtained using the semi-flexible method broken down by the 'Top Percent' parameter. These plots show there is no relationship between RMSD and pKi when the pose is selected by RMR6.

**Table S4:** Default parameters for CANDOCK.

| OPTION NAME | DEFAULT | DESCRIPTION |
|---|---|---|
| **NCPU** | -1 | Number of CPUs to use concurrently (use-1 to use all CPUs) |
| **RECEPTOR** | receptor.pdb | Receptor filename |
| **LIGAND** | ligands.mol2 | Ligand filename-c [ --config ] arg Configuration File |
| **SEEDS** | seeds.txt | Read unique seeds from this file, if it exists, and append new unique seeds if found |
| **MAX_NUM_LIGANDS** | 10 | Maximum number of ligands to read in one chunk |
| **SEEDS_PDB** | seeds.pdb | File to save full seeds into. |
| **PREP** | prepared_ligands.pdb | Prepared small molecule(s) are outputted to this filename |
| **CLUSTERFILE** | clustered_seeds.txt | Clustered representative docked-seed conformations output file |
| **NUM_UNIVEC** | 256 | Number of unit vectors evenly distributed on a sphere for conformation generation |
| **CLUS_RAD** | 2 | Cluster radius for docked seeds |
| **EXCLUDED** | 0.8 | Excluded radius |
| **CONF_SPIN** | 10 | Spin degrees for conformation generation |
| **MAX_FRAG_RADIUS** | 16 | Maximum fragment radius for creating the initial rotamers |

| | | |
|---|---|---|
| **GRIDPDB_HCP** | gridpdb_hcp.pdb | Grid pdb hcp file for output |
| **INTERATOMIC** | 8 | Maximum interatomic distance |
| **TOP_SEEDS_DIR** | top_seeds | Directory for saving top docked seeds |
| **GRID** | 0.375 | Grid spacing |
| **REF** | mean | Normalization method for the reference state ('mean' is averaged over all atomtype pairs, whereas 'cumulative' is a summation for atom type pairs) |
| **FUNC** | radial | Function for calculating scores 'radial' or 'normalized_frequency' |
| **SCALE** | 10 | Scale non-bonded forces and energy for knowledge-based potential [0.0-1000.0] |
| **POTENTIAL_FILE** | potentials.txt | Output file for potentials and derivatives |
| **DIST** | data/ csd_ complete_ distance_ distributions.txt | Select one of the interatomic distance distribution file(s) provided with thisprogram |
| **STEP** | 0.01 | Step for spline generation of non-bonded knowledge-based potential [0.0-1.0] |
| **COMP** | reduced | Atom types used in calculating reference state 'reduced' or 'complete'('reduced' includes only those atom types present in the specified |

| | | |
|---|---|---|
| | | receptor and small molecule, whereas 'complete' includes all atom types) |
| **CUTOFF** | 6 | Cutoff length [4-15]. |
| **UPDATE_FREQ** | 10 | Update non-bond frequency |
| **MAX_ITER** | 10 | Maximum iterations for minimization during linking |
| **FFTYPE** | kb | Forcefield to use 'kb' (knowledge-based), 'phy' (physics-based), or 'none' (do not calculate intermolecular forces) |
| **OBJ_DIR** | obj | Output directory for objective function. Setting this value will causethe KB potential to be read from disk. An empty string causes the objective function to be recalculated. |
| **TEMPERATURE** | 300 | Temperature to run the dynamic simulation at. |
| **MAX_ITER_FINAL** | 100 | Maximum iterations for final minimization |
| **MINI_TOL** | 0.0001 | Minimization tolerance |
| **AMBER_XML** | data/amber10.xml | Receptor XML parameters (and topology) input file |
| **WATER_XML** | data/tip3p.xml | Water XML parameters (and topology) input file |

| | | |
|---|---|---|
| **GAFF_DAT** | data/gaff.dat | Gaff DAT forcefield input file |
| **GAFF_HEME** | None | Gaff DAT file to use for Heme groups |
| **DYNAMIC_STEPS** | 1000 | Number of steps to do a dynamic simulation for. |
| **POS_TOL** | 1E-11 | Minimization position tolerance in Angstroms - only for KB |
| **DYNAMIC_STEP_SIZE** | 2 | Step size (in fempto seconds) |
| **INTEGRATOR** | verlet | Which integrator to use. Options are 'verlet', 'langevin', or 'brownian' |
| **GAFF_XML** | data/gaff.xml | Gaff XML forcefield and ligand topologyoutput file |
| **FRICTION** | 91 | Friction/Collision frequency for a dynamics simulation in 1/ps |
| **UPPER_TOL_SEED_DIST** | 2 | Upper tolerance on seed distance for getting initial conformations of dockedfragments |
| **TOL_SEED_DIST** | 2 | Tolerance on seed distance in-between linking |
| **ITERATIVE** | 1 (Implicit) | (=false) Enable iterative minimization during linking |
| **MAX_NUM_POSSIBLES** | 200000 | Maximum number of possibles conformations considered for clustering |

| | | |
|---|---|---|
| **CLASH_COEFF** | 0.75 | Clash coefficient for determining whether two atoms clash by eq. dist12 s< C * (vdw1 + vdw2) |
| **SPIN** | 60 | Spin degrees to rotate ligand. Allowed values are 5, 10, 15, 20, 30, 60, 90 |
| **MAX_POSSIBLE_CONF** | -1 | Maximum number of possible conformations to link (-1 means unlimited) |
| **DOCKED_DIR** | docked | Docked ligands output directory |
| **RMSD_CRYSTAL** | 1 (Implicit) | (=false) If the crystal ligand's pose was given,calculate RMSDs for each pose |
| **LINK_ITER** | 1000 | Maximum iterations for linking procedure |
| **DOCKED_CLUS_RAD** | 2 | Cluster radius between docked ligand conformations |
| **JIGGLE_SEED** | 1597463007 | Seed to use for randomization of top_seed positions. -1 will set the seed by random device, -2 will disable jiggle. |
| **MAX_ALLOW_ENERGY** | 0 | Maximum allowed energy for seed conformations |
| **TOP_PERCENT** | 0.05 | Top percent of each docked seed to extend to full molecule |
| **LOWER_TOL_SEED_DIST** | 2 | Lower tolerance on seed distance for getting initial conformations of dockedfragments |

| | | |
|---|---|---|
| **MAX_CLIQUE_SIZE** | 3 | Maximum clique size for initial partialconformations generation |
